# High Sensitivity and Low-Cost Flavin luciferase (FLUX)-based Reporter Gene for Mammalian Cell Expression

**DOI:** 10.1101/2021.07.04.451075

**Authors:** Jittima Phonbuppha, Ruchanok Tinikul, Yoshihiro Ohmiya, Pimchai Chaiyen

## Abstract

Luciferase-based gene reporters generating bioluminescence signals are important tools for biomedical research. Amongst the luciferases, flavin-dependent enzymes use the most common, and thus most economical chemicals. However, their applications in mammalian cells are limited due to their low signals compared to other systems. Here, we constructed <u>F</u>lavin <u>Lu</u>ciferase for Mammalian <u>C</u>ell Expression (FLUX) by engineering luciferase from *Vibrio campbellii* (the most thermostable bacterial luciferase reported to date) and optimizing its expression and reporter assays in mammalian cells. We found that the FLUX reporter gene can be overexpressed in various cell lines and showed outstanding signal-to-background in HepG2 cells, significantly higher than that of firefly luciferase (Fluc). The combined use of FLUX/Fluc as target/control vectors gave the most stable signals, better than the standard set of Fluc(target)/Rluc(control). We demonstrated that FLUX can be used for testing inhibitors of the NF-κB signaling pathway, validating FLUX applications for various assays in the future.

## INTRODUCTION

High throughput screening technology is important for biomedical research because it can be employed to find drug candidates against infectious and non-communicable diseases (NCDs). NCDs are chronic diseases not transmissible between people and have become the world’s major killers, accounting for 71% of all deaths globally (WorldHealthOrganization, 2021). Despite the rapid developments in screening technology which enable new drugs or active pharmaceutical ingredients to be discovered, such resources are not widely accessible around the world, and are especially lacking in developing countries due to the high costs. This has resulted in the majority of low- to middle-income countries (LMIC) to be unable to afford independent drug development programs, contributing to high numbers of premature deaths, mainly in the South-East Asian, Eastern Mediterranean and African regions (Martinez et al., 2020). Over 80% of the overall deaths in LMIC regions are caused by cardiovascular, (lung, esophagus, stomach and liver) cancers, chronic respiratory, congenital birth defects and digestive diseases (Gelband et al., 2015; Martinez et al., 2020). Therefore, development of effective and affordable high throughput assays or screening tools would allow researchers with limited funding to access similar technologies without such a high cost barrier, democratizing and distributing sustainable scientific development across the globe.

Reporter genes are among the common tools used for drug and bioactive compound screening. The technique can be used for monitoring cellular events associated with signal transduction, gene expression as well as disease progression (Naylor, 1999; Roda et al., 2004). Reporter genes can generate various signals for detection including absorbance, fluorescence, and luminescence. Among these detection methods, luminescence is the most sensitive techniques, providing detection limit as low as 10^−18^ – 10^−21^ moles of analytes. (Díaz-García & Badía-Laíño, 2019; Roda et al., 2004). Unlike fluorescence detection which requires excitation light that can create high background, bioluminescence does not require light input, thus eliminating photobleaching and lowering background signals. Bioluminescence usually generates a broad linear range of detection as well as high sensitivity (Roda et al., 2004; Thorne et al., 2010). It is widely used for monitoring cell progression, cytotoxicity, gene expression and cellular events relevant to a regulatory element, transcription factors and activities of bioactive compounds (Fan & Wood, 2007; Nakajima & Ohmiya, 2010; Riss et al., 2005; Thorne et al., 2010).

Bioluminescence is catalyzed by an enzyme class called luciferases, which is present in various organisms. Different luciferases use different substrates or luciferins with a common use of oxygen to generate light with specific wavelengths and signals. Current luciferin and luciferase systems include 1. coelenterazine substrates for *Renilla, Gaussia*, and *Oplophorus* luciferases, 2. D-luciferin substrate for firefly and Click beetle luciferases, 3. flavin-dependent system for bacterial luciferase, 4. cypridina luciferin-based system for *Cypridina* and *Porichthys* luciferases, 5. tetrapyrrole-based luciferins for luciferases from Dinoflagellates and Euphausiids 6. 3-hydroxyhispidin-based system for fungal luciferase (Fleiss & Sarkisyan, 2019). The first two groups of luciferins/luciferases are among the most commonly used systems as gene reporters because their light signals are high, giving high sensitivity in detection applications. Currently, the D-luciferin and firefly luciferase (Fluc) is the most widely used system which gives good signals and robustness for its applications. Although the coelenterazine system is simpler than other systems because it only requires one substrate plus oxygen, the compound is unstable and can release photons even in the absence of the *Renilla* luciferase (Rluc), resulting in relatively high background (Wang & El-Deiry, 2003; Zhao et al., 2004). The coelenterazine/*Renilla* luciferase system is also quite expensive; its price per assay is ∼1.4-times that of the D-luciferin/firefly luciferase. Tetrapyrrole-based system is highly active under acidic conditions, and the substrate is not commercially available, while the 3-hydroxyhispidin luciferase has just recently been discovered and still requires further investigation to fully understand its mechanisms. Among all existing luciferase systems, the flavin-dependent enzyme uses the simplest (thus most economical) substrates in which the price per assay would be 1/100 of the firefly system. However, its applications as a reporter gene in mammalian cells are limited due to its low signal. The successful development of the flavin-dependent luciferase as a gene reporter assay in mammalian cells would contribute to technology enabling high throughput screening tools that are ∼100-time and ∼150-time less expensive than the firefly and *Renilla* systems, respectively. In addition, the flavin-dependent luciferase with high bioluminescence signals can also work in complement with firefly luciferase as a dual reporter system typically used in mammalian cell expression experiments.

Bacterial luciferase (Lux) catalyzes bioconversion of reduced flavin, long chain aldehyde and oxygen to result in oxidized flavin, long chain carboxylic acids and H_2_O with concomitant light production with maximum emission around 490 nm. The enzyme consists of α and β subunits which are individually encoded by the genes *luxA* and *luxB*, respectively in the *lux* operon (Suadee et al., 2007). The *lux* operon also contains the *luxCDE* genes encoding multi-enzyme fatty acid reductase complex (LuxCDE) that converts fatty acid to aldehyde to supply the Lux reaction (Meighen, 1991). Several species of luminous bacteria also contain the luxG gene which encodes for a flavin reductase that catalyzes the production of reduced flavin, a substrate for Lux (Nijvipakul et al., 2008). Reduced flavin is the first substrate to bind to Lux, followed by the oxygen reaction to generate the C4a-peroxyflavin intermediate which attacks an aldehyde substrate and generates the following C4a-peroxyhemiacetal intermediate. The cleavage of the O-O bond in C4a-peroxyhemiacetal yields fatty acid and the excited C4a-hydroxyflavin, which emits the blue-green light with λ_max_ around 490 nm (Suadee et al., 2007; Tinikul & Chaiyen, 2014; Tinikul et al., 2020). In the past, several studies have investigated the expression of Lux in mammalian cells, but most of them only investigated Lux from the terrestrial microbe, *Photorhabus luminescens* (*Pl*_Lux) (Class et al., 2015; Close et al., 2010; Cui et al., 2014; Eldridge et al., 2007; Gregor et al., 2019; Patterson et al., 2005; Sanseverino et al., 2005; Xu, Close, Webb, Price, et al., 2013; Xu, Close, Webb, Ripp, et al., 2013; Xu et al., 2015; Xu et al., 2014). The results of these studies showed that mammalian cells can overexpress *Pl*_Lux, but their signals are rather low (Close et al., 2010; Cui et al., 2014; Gregor et al., 2019; Patterson et al., 2005; Tehrani et al., 2014). To the best of our knowledge, none of the studies has compared the analytical power of *Pl*_Lux to other commonly used luciferases side-by-side nor demonstrated that the *Pl*_Lux can be used in substitution of firefly or *Renilla* luciferases as a gene reporter.

We proposed that Lux from *Vibrio campbellii* (*Vc*_Lux) is another attractive system to be used as a gene reporter. *Vc*_Lux is the most thermostable bacterial luciferase reported to date and when expressed in *E. coli*, can generate about 100-fold brighter light than the enzyme from *Vibrio harveyi* (*Vh*_Lux) (Suadee et al., 2007; Tinikul & Chaiyen, 2014; Tinikul et al., 2013; Tinikul et al., 2012). As *Vh*_Lux generates light 5-fold brighter than that of *Pl*_Lux (Westerlund-Karlsson et al., 2002), it can be assumed that light generated by *Vc*_Lux is much brighter than that of *Pl*_Lux. Although *Vc*_Lux with two subunits fused *via* a linker can be overexpressed in mammalian cells, the system previously constructed gives very low bioluminescent signals (Phonbuppha et al., 2021; Tinikul et al., 2012), making it still impractical for gene reporter applications in mammalian cell systems.

In this work, we improved the expression level of the fusion *Vc*_LuxAB (Lux) by optimizing its codon usage and modified redox and cell lysis reagents for enzyme assay cocktails. The newly developed <u>F</u>lavin <u>L</u>uciferase for Mammalian <u>E</u>xpression (FLUX) showed remarkable performance, yielding >400-times brighter signals than the system without optimization. These optimizations, for the first time, significantly increased the bioluminescence signals of the bacterial luciferase to be close to the firefly enzyme (only about 20-fold lower). Furthermore, the FLUX system has the added advantage of generating much lower background signal than Fluc or Rluc. We explored the use of FLUX in various cell-lines including HEK293T, NIH3T3, COS1, and HepG2 cells in comparison to Fluc. The results showed that FLUX can be expressed well in all four types of cell-lines and its signal-to-background ratio is even higher than FLuc in HepG2 cells. As transient transfection of gene reporters generally requires two different types of light signal generators, one as the target vector for addressing experimental effects and another as the control vector for evaluating the transfection efficiency, we thus explored the use of three luciferases (Fluc, Rluc and FLUX) in this combined luciferase-reporter gene assays. We found that the combined use of FLUX as the target vector and Fluc as the control vector gave the best result. This combined FLUX/Fluc luciferase-reporter gene assay was investigated for its sensitivity in investigating the effects of Tumor Necrosis Factor (TNF)-alpha and inhibitors on the NF-κB cell signaling pathway. The results showed that the FLUX/Fluc reporter system gave similar EC_50_ values compared to the use of Fluc/Rluc as target/control vectors, validating the potential use of FLUX in gene reporter applications. As luciferase reporter systems are widely used in the biomedical community (In year 2020 alone, more than 22,000 publications used luciferase reporters, (Figure 3-figure supplement 1, pink bars), the 1/100 price reduction of FLUX would make the system attractive as an alternative gene reporter with significant cost reduction, while maintaining good capability in high throughput screening applications in the future.

## RESULTS

### Optimization of a fusion bacterial luciferase (*lux*) gene expression in mammalian cells

We first constructed and optimized a fusion Lux from *Vibrio cambellii* in which both α and β subunits are linked *via* an artificial linker obtained from modification of the intergenic sequence linking luxA and luxB by adding the nucleotide (**G**) upstream of the stop codon of *lux*A gene (TAA) and mutating the start codon of luxB from ATG to **CAG** in order to abolish the stop codon and avoid any internal initiation of translation of *lux*B. The sequence of the artificial linker is **G**TAATTAATATTTTCGAAAAGGAAAGAGAC**CAG**, which encodes for 11 amino acids (VINIFEKERDQ). This allows Lux to be overexpressed mono-cistronically in mammalian cells (Tinikul et al., 2012). The previously published sequence of the *lux* gene from *Vibrio campbeiill* (*Vc_lux*) (Tinikul et al., 2012) was analyzed for the Codon Adaptation Index (CAI) using the GenScript Rara Codon Analysis Tool. The analysis showed that the original *lux* gene has a CAI value of only 0.67. We optimized the codon of the *lux* gene (designated as the *FLUX* gene) to obtain the CAI value of 0.88 (sequence shown in Figure 1-figure supplement 1 and deposited in NCBI database with GenBank number MZ393808) which should be suitable for expression in mammalian cells as a CAI value of higher than 0.8 is generally recommended for good gene expression. Expression of the *FLUX* gene using the pGL3 vector under a constitutive SV40 promoter was investigated in comparison with the original *lux* gene. Each system was also co-transformed with the pRL-TK vector to normalize transfection efficiency. The results showed that FLUX showed significant improvement in the bioluminescent signals, yielding a 414±33-fold increase in bioluminescent signals compared to that of the lux (Figure 1A). We further investigated whether this signal increment was due to higher levels of protein expression using western blot analysis. The western blot results showed that the protein overproduced by the *FLUX* gene was also significantly higher than that of *lux* gene (Figure 1B). Our data indicate that with suitable codon usage, the FLUX system can generate light reasonably well in mammalian cells.

**Figure 1.**
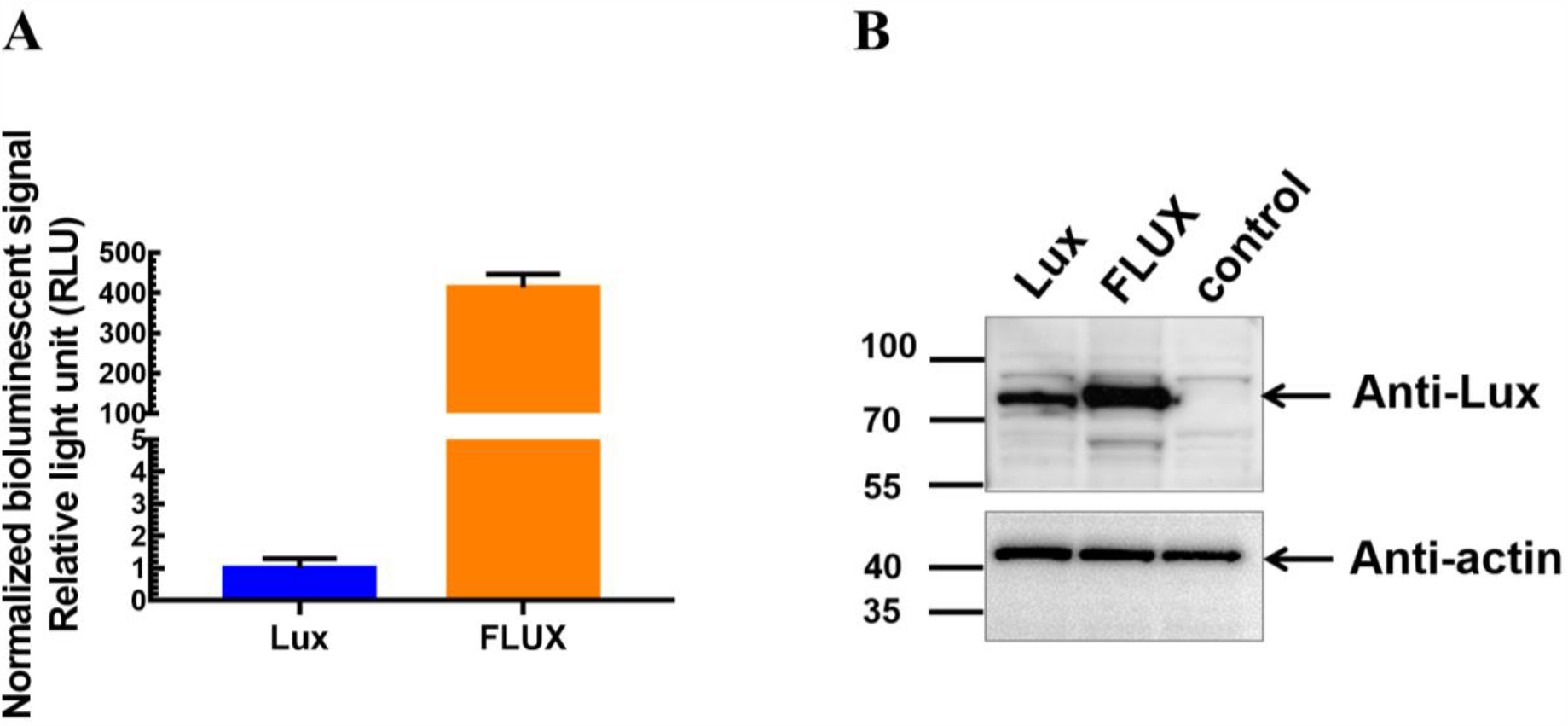
Expression of the original fusion bacterial luciferase gene (*lux*) and codon optimized (*FLUX*) in HEK293T cells. Vectors of pGL3 [*lux*/SV40] or pGL3[*FLUX*/SV40] (0.07 pmol each) and 0.007 pmol of pRL-TK vector (internal control) were co-transfected into HEK293T cells. After 48 hours. of transfection, cells were collected, and Lux Lysis Reagent (LLR) was added, and the protein expression and luciferase activity were measured. **A**. The activity of Lux was monitored by adding a solution (100 µL) of reagent cocktail consisting of 5 µM FMN, 100 µM HPA, 10 µM decanal and 100 µM NADH into a solution of cell lysate that was freshly mixed with 50 mU of reductase C_1_. The luminescence signal was monitored for 10 sec with a 2-sec delay using an AB-2250 single tube luminometer (ATTO Corporation, Japan). The *Rluc* activity was measured using the *Renilla* luciferase assay reagent [E2810, Promega Corporation) according to the manufacturer’s instructions. The Lux activity was normalized with the Rluc activity and reported as normalized bioluminescence signals by comparing the fold change in activities. Error bars indicate the standard deviation (n = 4). **B**. Expression level of each Lux was measured using western blot analysis with anti-flux antibody for detection of Lux protein overexpressed in cells containing the Lux (pGL3[*lux*/SV40]) and FLUX (pGL3[*FLUX*/SV40]). The control sample was lysate from cells without any transfection. Anti-actin was used for detection of actin as a housekeeping gene for normalizing the signals. Signal detection was carried out using secondary staining with HRP-conjugation and chemiluminescent reagents for HRP (EzWestLumi plus, ATTO Cooperation, Japan). The luminescence from HRP activity was imaged on a WSE-6100 LuminoGraph I Gel documentation system.

### Optimization of lysis and assay reagents for maximum light output

Lysis reagent is a buffer solution used for lysing cells and stabilizing proteins of interest. Ideally, lysis reagents should be mild, efficient in causing cell lysis and compatible with reagents used in downstream assays. The main additives in lysis reagents are mainly detergent, protease inhibitors and protein stabilizers in a suitable buffer (Leibly et al., 2012). Commercially available reagents include Reporter Lysis Buffer (RLB, Promega), Luciferase Cell Lysis Reagent (CCLR, Promega), Passive Lysis Buffer (PLB, Promega). The first two groups of lysis reagent are suitable for Fluc, while PLB is a special reagent that can be used for both Fluc and Rluc (Sherf et al., 1996). In this work, we developed a lysis reagent for the FLUX system based on common reagents available in the laboratory.

We aimed to find suitable reagents including lysis detergent, protease inhibitor and protein stabilizing agents for the FLUX assays. First, Triton X-100 and CHAPS, which are nonionic and zwitterionic amphipathic compounds, respectively were used to lyse cells by solubilizing lipids and protein in the membrane and creating pores within the membrane for full cell lysis. Their effects on FLUX activity were investigated by measuring the bioluminescent signals of the purified recombinant Lux in various detergent concentrations. The results showed that Lux activity was susceptible to TritonX-100 because even the lowest TritonX-100 concentration (0.125 % w/v) resulted in a 10% decrease in activity of Lux, while 0.5 % (w/v) TritonX-100 decreased the activity by 50 % (Figure 2A). The Lux activity is more stable in the presence of CHAPS, demonstrating stability in CHAPS concentrations ranging from 0.63 up to 2.5 % (w/v), while 5 % (w/v) CHAPS resulted in a drop in activity of around 50% (Figure 2B). These results suggest that 1-2 %(w/v) of CHAPS is suitable for lysing the mammalian cells transfected with the *FLUX* reporter gene. For protease inhibitors, EDTA which is a metal chelator capable of chelating metal cofactors of several metalloproteases generally produced in mammalian cells was chosen as a protease inhibitor additive. We investigated the effects of EDTA on FLUX activity by measuring bioluminescence signals of the purified recombinant Lux in various concentrations of EDTA. The results showed that EDTA does not affect FLUX activity; the system retained nearly 100% of the bioluminescence in 0.25 - 5 mM EDTA (Figure 2C).

**Figure 2.**
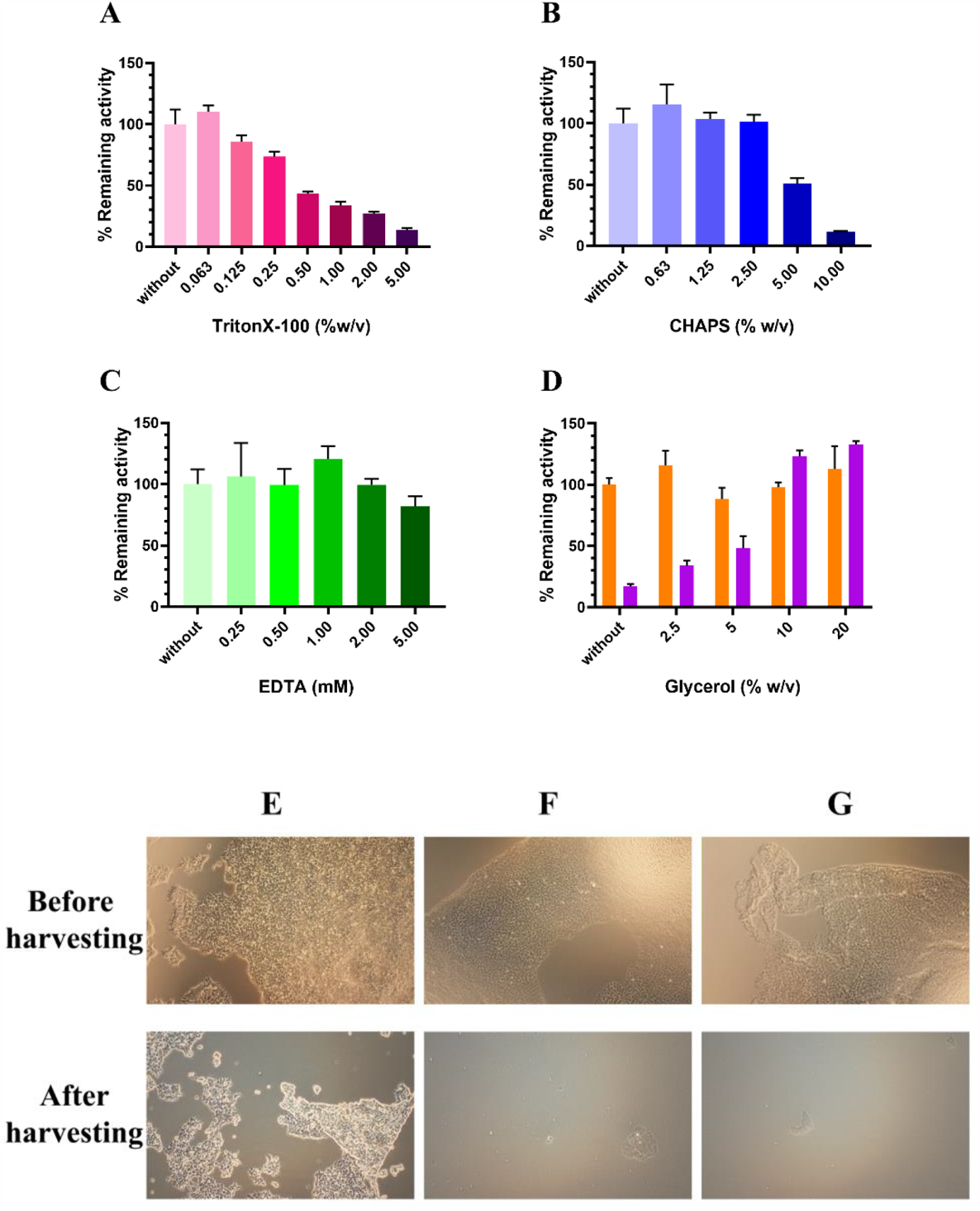
Effects of various additive compounds on the purified Lux activity. **A**. Triton-X100, **B**. CHAPS, **C**. EDTA, and **D**. Glycerol before (orange) and after three freeze-thaw cycles (purple). Efficiency comparison of cell harvesting using different lysis reagents including **E**. (50 mM sodium phosphate buffer pH 7.0), **F. (**Lux Lysis Reagent (LLR), this study) and **G**. Cell Culture Lysis Reagent (CCLR, Promega). **A-D**, The effects of each additive on Lux activity was investigated by pre-mixing the purified Lux solution with each additive at various concentrations and incubating for 15 min before measuring Lux activity compared to purified Lux. The activity of Lux was monitored by adding a solution (100 µL) of assay cocktail consisting of 5 µM FMN, 100 µM HPA, 10 µM decanal and 100 µM NADH into a solution of cell lysate that was freshly mixed with 50 mU of reductase C1. The luminescence signal was monitored for 10 sec with a 2-sec delay using an AB-2250 single tube luminometer (ATTO Corporation, Japan). Each Lux activity was divided by the Lux activity measured in the absence of any additive compounds to obtain a remaining activity. The remaining activity was multiplied by 100 to obtain %remaining activity. Error bars indicate standard deviations (n = 4). Effects of various lysis reagents on cell detachment were investigated by plating 1×10^5^ HEK293T cells on a 24-well plate for 24 hours before adding 100 µL of lysis reagent and then monitoring HEK293T cell morphology using an IX73 inverted light microscope (Olympus). Cells were incubated at room temperature with rocking for 15 min before collecting the detached cells and the remaining HEK293T cells on the culture plate were examined using the IX73 inverted light microscope (Olympus).

For an additive to stabilize proteins during freeze-thaw processes, we chose to explore the use of glycerol for this purpose. Because the activity assays usually cannot be performed on the same day as the cell harvesting and lysis process, cell lysates are generally kept frozen before thawing later for activity measurement. As glycerol is a common protein stabilizer which can prevent ice crystallization which destroys protein structures (Dashnau et al., 2006), the effects of glycerol on Lux activity were investigated during freezing-thaw processes by measuring the bioluminescence signals of Lux before and after three freeze-thaw cycles. The results showed that glycerol (2.5 to 20 % (w/v)) had no effect on Lux signals because it could retain nearly 100% of the Lux activity over the entire range of glycerol concentrations investigated (Figure 4D, Orange). However, only 10-20% (w/v) of glycerol allowed retention of ∼100% of Lux activity after three freeze-thaw cycles (Figure 4D, Purple). These results suggest that at least 10 % (w/v) glycerol is suitable for stabilizing the protein during the freezing-thawing process.

A final formula of lysis reagents for the FLUX reporter gene or Lux Lysis Regent (LLR) developed in this work consists of 1 % (w/v) CHAPS, 1 mM EDTA, 10 % (w/v) glycerol in 50 mM sodium phosphate buffer pH 7.0 which is typically used in the assay reaction. Thus, we further investigated the efficiency of LLR compared to a buffer without any additive reagent by monitoring cell morphology on the culture plate before and after harvesting the cells. Commercial lysis buffers for harvesting adherent cells such as Cell Culture Lysis Reagent (CCLR, Promega) was also compared to evaluate the efficiency of LLR. Results showed that most of the adherent cells could be harvested by LLR (Figure 3F, after harvesting), while using a buffer without any additives resulted in most of the cells remaining adhered to the culture plate (Figure 3E, after harvesting). The LLR harvesting efficiency is comparable to that of CCLR (Figure 3G, after harvesting), suggesting that LLR is a suitable lysis buffer to remove and lyse adherent cells.

**Figure 3.**
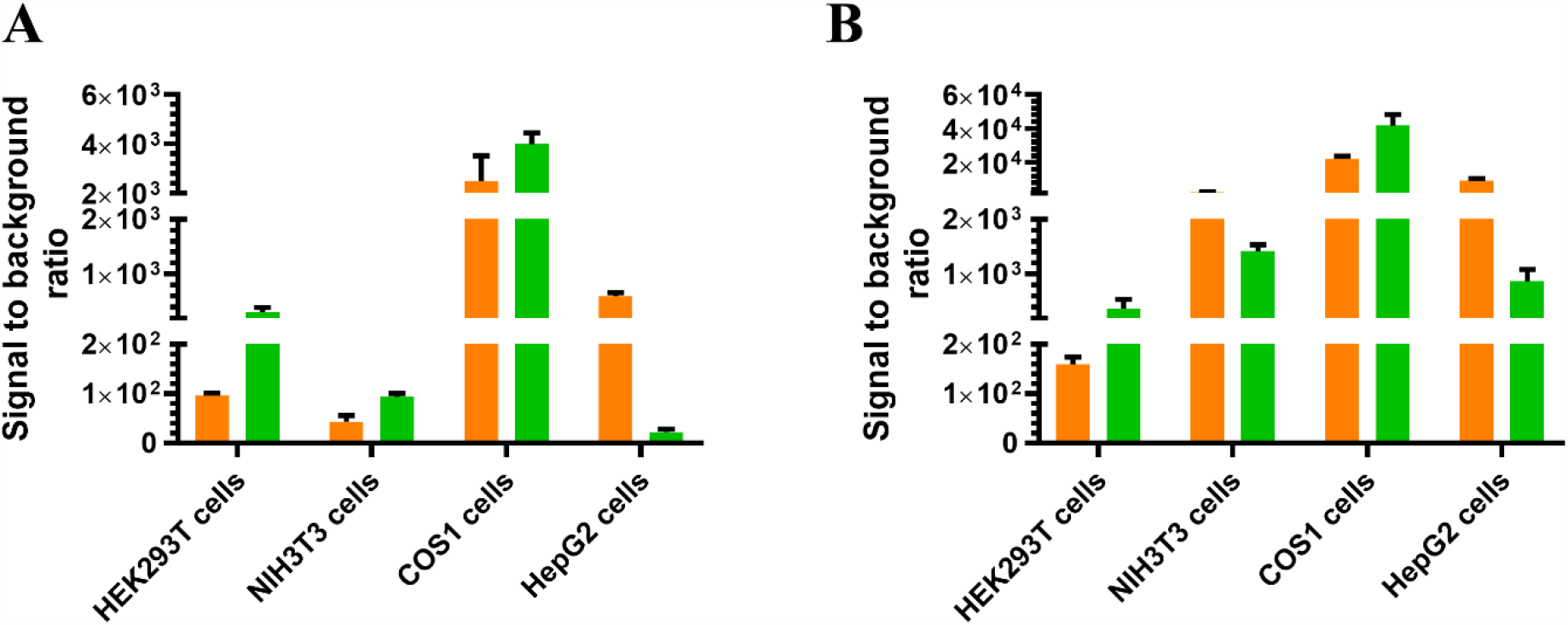
Comparison of signal to background ratios of FLUX (orange) and Fluc (green) gene reporter systems **A**. Systems constructed in the pGL3 backbone vector and **B**. Systems constructed in the pGL4 backbone vector. Various cell types including HEK293T, NIH3T3, COS1, and HepG2 cells were used for testing bioluminescence signals. HEK293T, NIH3T3, COS1, and HepG2 cells were co-transfected with each reporter gene (0.07 pmol) and the internal control pRL-TK vector (0.007 pmol). Cells were collected at 48-hours after transfection by washing the culture with Passive Lysis Buffer or Lux Lysis Reagent (LLR). Then, luciferase activity was measured by adding a 100 µL of assay cocktail consisting of 5 µM FMN, 100 µM HPA, 10 µM decanal and 100 µM NADH into a solution of cell lysate that freshly mixed with 50 mU of reductase C_1_. The luminescence signal was monitored for 10 sec with a 2-sec delay using an AB-2250 single tube luminometer (ATTO Corporation, Japan). The Fluc and Rluc activities were measured using firefly luciferase and *Renilla* Luciferase Assay reagents, respectively [E1500 and E2810, Promega Corporation) according to the manufacturer’s instructions. The luciferase activity under the constitutive SV40 promoter was divided by their Rluc activity to obtain the normalized luciferase activity. The normalized signal of SV40 promoter vector was divided by the normalized signal from the corresponding promoterless vector to obtain a value of signal to background ratio. Error bars indicate the standard deviation values from experiments with n = 4.

**Figure 4.**
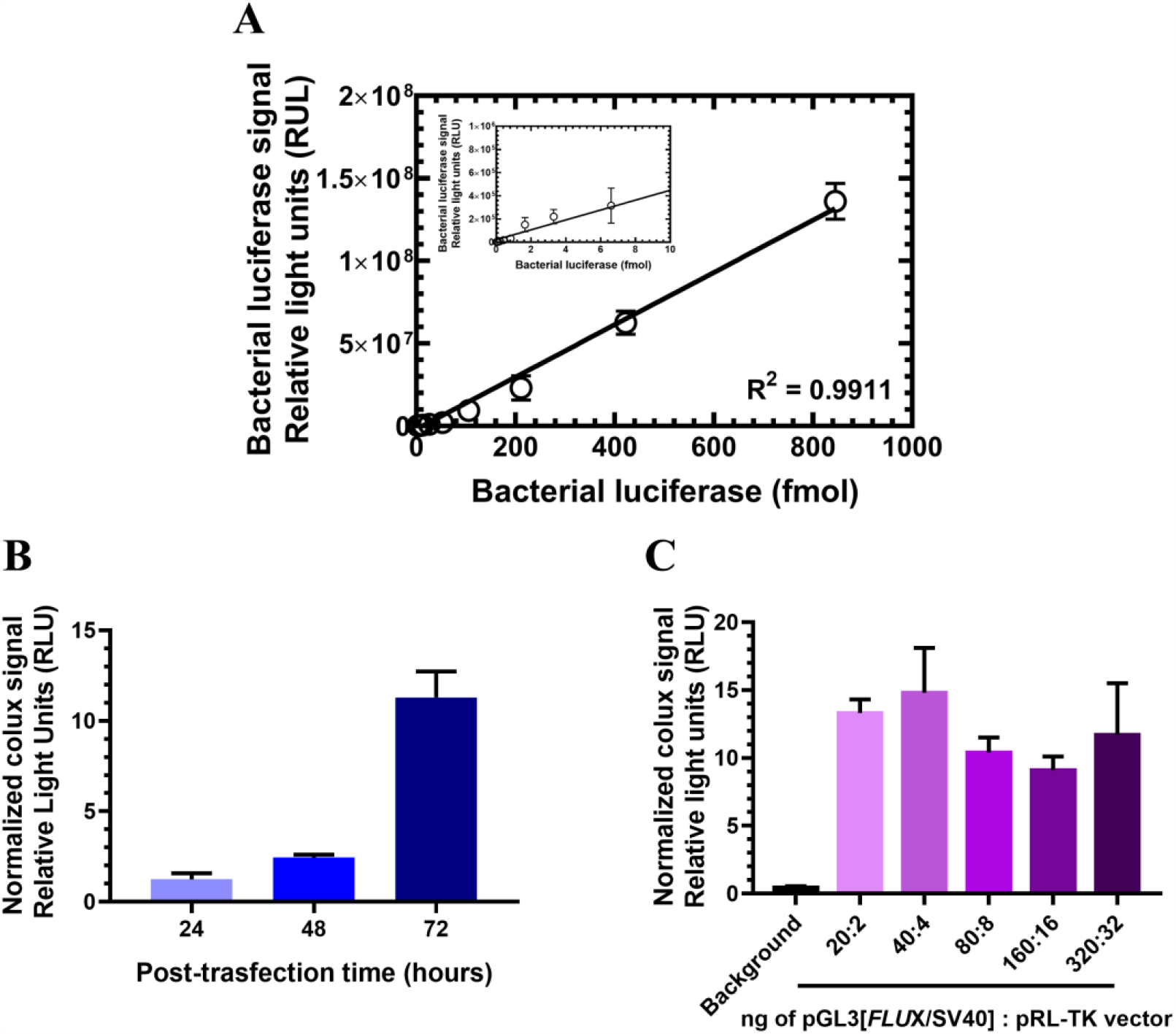
FLUX signals and their influencing factors. **A**. A correlation of responsive signals and the amount of purified recombinant Lux. 2-Fold dilutions of Lux were made from 850 fmoles to 25 amoles for determining a correlation between Lux signal at various amount of the enzyme. The correlation of responsive signals with amount of Lux was plotted and shown in the insert figure. The coefficient of determination (R^2^) was analyzed by GraphPad Prism Version 8 software shown in the figure. **B**. Effects of a post-transfection period on FLUX signals. HEK293T cells were transfected with the pGL3 [*FLUX*/SV40] vector (0.07 pmol) and 0.007 pmol of the internal control pRL-TK vector. Cells were collected at 24-, 48-, and 72-hours post-transfection using Lux Lysis Reagent (LLR) and the resulting luciferase activities were independently measured. **C**. Normalized FLUX signals at various amounts of FLUX vector. A pGL3[*FLUX*/SV40] vector was co-transfected with pRL-TK vector as internal control vector at target:control vector amount of 20:2, 40:4, 80:8, 160:16, and 320:32 ng into HEK293T cells. The activity of Lux was monitored by adding a 100 µL of assay cocktail consisting of 5 µM FMN, 100 µM HPA, 10 µM decanal, and 100 µM NADH into a solution of cell lysate that was freshly mixed with 50 mU of reductase C_1_. The luminescence signal was monitored for 10 sec with a 2-sec delay using an AB-2250 single tube luminometer (ATTO Corporation, Japan). The Rluc activity was measured using *Renilla* Luciferase Assay Reagent [E2810, Promega Corporation] according to the manufacturer’s instructions. The FLUX activity was divided by their Rluc luciferase activity to obtain normalized luciferase signals. Error bars indicate the standard deviation values (n = 4).

As our data showed that LLR is a suitable lysis reagent to harvest adherent cells and help stabilize the Lux protein, we further applied LLR for lysing adherent cells expressing FLUX by measuring the FLUX activity. Previously, LUX activity could be assayed using the reagents consisting of 10 µM FMN, 200 µM HPA, 200 µM NADH, and 20 µM decanal in 50 mM sodium phosphate buffer pH 7.0. Typically, 20-50 µL of cell lysate was mixed with 50 mU reductase C_1_ before adding the assay cocktail to monitor bioluminescent signals (Tinikul et al., 2012). In this work, we found that when using LLR for cell harvesting and lysis, cell lysate volume could be reduced by 10-fold, possibly because LLR better lyses adherent cells and stabilizes the Lux protein (Data not shown). As the cell lysate amount required was reduced, we further optimized the cocktail assay reagents to minimize the cost, while maintaining high bioluminescent signals. We found that the Lux assay cocktail consisting of 5 µM FMN, 100 µM HPA, 100 µM NADH, and 10 µM decanal in 50 mM sodium phosphate buffer pH 7.0 provides bioluminescent signals comparable to the previously used Lux cocktail (data not shown) which contained the same substrates with most at two-fold higher concentrations and thus would be twice as expensive. Altogether, our results indicate that the optimization of lysis and assay reagents could maximize bioluminescent output and minimize the FLUX reporter gene’s assay cost.

### Comparison of light generated from FLUX and Fluc

To evaluate whether the FLUX system can be used as a gene reporter in transient transfection for probing molecular events in mammalian cells, we carried out experiments to measure light signal to background (S/N) ratios to compare the sensitivity of FLUX to the Fluc reporter gene, which is broadly used in a wide variety of applications. Bioluminescence readout signals generally contain target signals and unspecific signals from background, stray light and detector offset, etc. (Alkemade et al., 1978). Therefore, a S/N ratio, not an absolute signal value, is generally used as a parameter to represent measurement signals. Generally, signal can be measured from bioluminescence generated under the control of a constitutive or inducible promoter, while background can be measured from bioluminescence generated in the absence of any activating element (Paguio et al., 2005). The S/N value can evaluate the strength of the promoter or measure the potency of compounds/metabolites under investigation such as inhibitors or inducers of the promoter (Wunsch et al., 2005). For comparison and benchmarking, we chose two vector systems commonly used for expression of the Fluc reporter gene, pGL3[*luc*+] and pGL4[*luc*2] vectors (Promega, USA), to construct the expression vector of FLUX to compare with the bioluminescence signals of Fluc. The pGL4[*luc*2] vector is the newest series of Fluc reporter genes with a significantly improved signal to background ratio compared to the previous version pGL3[*luc+*] vector (Paguio et al., 2005). However, the pGL3[*luc+*] vector is still widely used in biological research, with 5040 published articles in 2020 (Figure 3-figure supplement 1, green bars). Therefore, the *FLUX* gene was inserted into the pGL3 and pGL4 vectors downstream of the constitutive SV40 promoter as well as in the same position in a vector without SV40 promoter to construct vector sets with and without SV40 (Figure 3-figure supplement 2). All systems were independently co-transfected with the vector containing Rluc as an internal control vector in various cell types including HKE293T, NIH3T3, COS1, and HepG2 cells.

The results showed that the FLUX system generated less light intensity both in target and background signals than Fluc in both pGL3[*luc+*] and pGL4 [*luc2*] vectors (Figure 3-figure supplement 3). This is due to the nature of the low quantum yield of Lux compared to Fluc (Hastings et al., 1965; Niwa et al., 2010). However, analysis of signal to background ratios (S/N) showed that S/N values of FLUX and Fluc expressed in the pGL3 expression vector are comparable in NIH3T3 and COS1 cells (Figure 3A). S/N values of FLUX are less than the Fluc in HEK293T cells while they are considerably higher than Fluc in HepG2 cells (Figure 3A). These S/N ratio behaviors are similar to the experiments comparing the FLUX and Fluc expression using the pGL4 vector (Figure 3B). The higher S/N ratio of FLUX in HepG2 cells than Fluc is caused by the higher background signals of Fluc than FLUX, whereas the two systems generated comparable target signals (Figure 3-figure supplement 3). The higher background signals in the Fluc system (in both *luc*+ and *luc*2 genes) might be the result of anomalous expression of the reporter gene caused by cryptic regular-binging site or/and enhancer element (Paguio et al., 2005). With the origin of FLUX being from bacteria, such anomaly is less evident in mammalian systems.

These data in Figure 3 clearly suggest that FLUX functioned well as gene reporters in HepG2 and COS1 cells because the S/N values of the FLUX systems were quite high in both cells with both vector types. For the NIH3T3 cells, the FLUX system also showed higher S/N values than the Fluc system with the pGL4 vector, also implying that the FLUX system should be able to serve as a gene reporter in NIH3T3 as well. Among all cell types, the FLUX system gave lower S/N ratio than the Fluc system in HEK293T cells (about 2-folds). Based on these data in Figure 3 alone, the ability of FLUX to serve as a gene reporter system in HEK293T cells was questionable. We thus investigated the ability of FLUX to serve as a gene reporter in HEK293T cells in the following sections. We chose to validate the function of FLUX in HEK293T cells to prove the functions of FLUX in the least favorable expression systems. If FLUX can be used as a gene reporter in HEK293T, the results inevitably endorse the use of FLUX in more favorable HepG2, COS1 and NIH3T3 cells.

### Exploring capability and robustness of the FLUX gene reporter system

To evaluate the use of FLUX system as a gene reporter, we explored the range of linear detection for FLUX to evaluate the sensitivity and working range of the system, and also investigated parameters that can affect the transient transfection process including cell growth period and amount of vector used for transfection.

First, the sensitivity and linear range of detection by FLUX were investigated by measuring the bioluminescence signals generated by various amount of the purified Lux. The results showed a broad linearity range with at least eight orders of detection magnitude; this is equivalent to a broad dynamic range of 0.25 fmoles to 850 fmoles (Figure 4A). These data indicate that bioluminescent signals directly depend on the amount of Lux over a wide range, also enabling limits of detection down to a few molecules at amole levels (10^−18^ mole). The wide range of FLUX detection limit (eight orders of magnitude) is similar to the detection range of Fluc and is wider than that of Rluc (about seven orders of magnitude) (Sherf et al., 1996). However, Fluc and Rluc give higher sensitivity than FLUX because their limit of detection (LoD) is lower. The LoDs of Fluc, Rluc and Lux are 0.01, 0.3 and 25 amoles of enzymes, respectively which correlate with their quantum yields (0.48, 0.05, and 0.16 for Fluc, Rluc, and Lux, respectively) and their catalytic turnovers (1.6 s^-1^, 1.9 s^-1^, and 0.005 s^-1^, for Fluc, Rluc, and Lux, respectively) (Branchini et al., 1998; Lei et al., 2004; Loening et al., 2006; Matthews et al., 1977; Niwa et al., 2010; Sherf et al., 1996; Suadee et al., 2007; Sucharitakul et al., 2007). However, it should be noted that this LoD level was based on the purified enzymes which is not directly relevant to the FLUX expression in mammalian cells. We later showed that transfection of 10^5^ cells, a level typically used in biological research with 20 ng of the FLUX vector can yield good signals for practical experiments (Figure 4-figure supplement 2 and see more results below).

Second, we investigated signals generated by FLUX after transfection. Transfected cells are typically harvested for analyzing levels of gene expression 24 hours and up to 72 hours post-transfection. A suitable time for cell harvesting is different, depending on cell type, research goals and specific expression characteristics (Doyle & Promega, 1996). Therefore, normalized FLUX activities in HEK293T cells transfected with the pGL3[FLUX/SV40] vector and the pRL-TK vector as an internal control at various time points after transfection were measured. Results showed that the luciferase activities in cell lysate increased over time (Figure 4B). The FLUX signal at 24-hours post-transfection was significantly higher than the non-transfected cells by at least 3 orders of detection magnitude and was much higher than the background signal (Figure 4-figure supplement 1, orange). The bioluminescent signals at 48- and 72-hours post-transfection were two-fold and nine-fold higher than that at 24-hours post-transfection, respectively (Figure 4B). The results suggested that although the signal from 24-hours post-transfection was sufficient for typical measurement, a longer period such as 48 or 72-hours could increase the signal significantly. The higher signals upon longer post-transfection times indicate that Lux is stable inside the cells and could accumulate signals upon prolonged cell growth. The stability of the intracellular FLUX protein makes the system suitable for high-throughput screening applications.

Third, we explored the suitable amount of vector required for transfection. Typically, a 96-well culture plate is commonly used for high-throughput screening applications and the amount of vector of 100 ng is normally recommended for transfection per well. We investigated the effects of the vector amount used for transfection on the Flux signals using FLUX/Rluc combination as a model for investigation. The amount of FLUX and Rluc reporter genes was varied from 20:2 ng to 320:32 ng, covering a range recommended for transfection and maintaining a final ratio of 10:1. The transfected cells were harvested at 24-hours post-transfection and the bioluminescent signals were measured in cell lysate. The results showed that even at the lowest amount of the pGL3[FLUX/SV40] vector (20 ng), a high S/N ratio could be obtained (Figure 4-figure supplement 2, orange,). Both FLUX and Rluc signals could be increased directly according to the amount of transfected vector (Figure 4-figure supplement 2, orange, and purple, respectively). Results in Figure 4C showed that the normalized FLUX signals which were obtained from dividing the original FLUX with Rluc signals were similar at all ratios investigated. The data suggest that FLUX could be used as a target reporter gene with requirement of only 20 ng per transfection.

### Combined use of FLUX, Fluc and Rluc as target/control vectors for luciferase-reporter gene assays

In transient transfection experiments, two types of vectors including target and control vectors are generally required, and the results shown in the previous section indicate that the FLUX system developed here gives signals suitable for reporter gene applications. We thus investigated the combined use of FLUX *versus* Fluc and Rluc in target/control reporter sets including 1. FLUX/Rluc, 2. FLUX/Fluc and 3. Fluc/FLUX in comparison with 4. the standard Fluc/RLuc combination set generally used in luciferase bioreporter experiments. These combined vector sets with varying ratios of target:control vectors from 10:1 to 10:20 were used for measuring their light signals. Ideally, consistency of light signals with varying ratios of vectors would indicate robustness of a reporter system as a measurement tool. In real practical experiments, concentrations of target and control vectors may not be exactly the same at each transfection reaction and a good reporter tool should not be overly affected by this variation as this could create artifact signals that interfere with the effects of the experimental parameters one wants to address (Schagat et al., 2007). In this experiment, we measured signals generated by target reporters with varying amounts of control reporters (Figure 5-figure supplement 1) and compared these values with the target reporter signals in the absence of control vector (calculated as % signal measured, Figure 5). Results in Figure 5 showed that the combination set 2. FLUX/Fluc, showed the most consistent light signals when varying the vector ratios because their percent signal measured was almost unperturbed even at the highest amount of control vector (target:control = 10:20) (Figure 5B). The use of FLUX/Rluc is also acceptable for target:control vector ratios of up to 10:2. In the standard combination of Fluc/Rluc (Figure 5D) and FLUX/Rluc (Figure 5A), the systems gave consistent percent signal measured with target:control ratios of 10:1-10:2, and the signals decreased at the higher target:control ratios. The behavior of the standard Fluc/Rluc combination were similar to those reported previously (Schagat et al., 2007). The results also showed that the combined use of Fluc/FLUX gave the highest variation in percent signal measured when the target:control vector ratios were varied (Figure 5C). Altogether, the data indicate that the combined use of FLUX/Fluc (Set 2) gave the most ideal properties for their combined use as a target and control reporter set.

**Figure 5.**
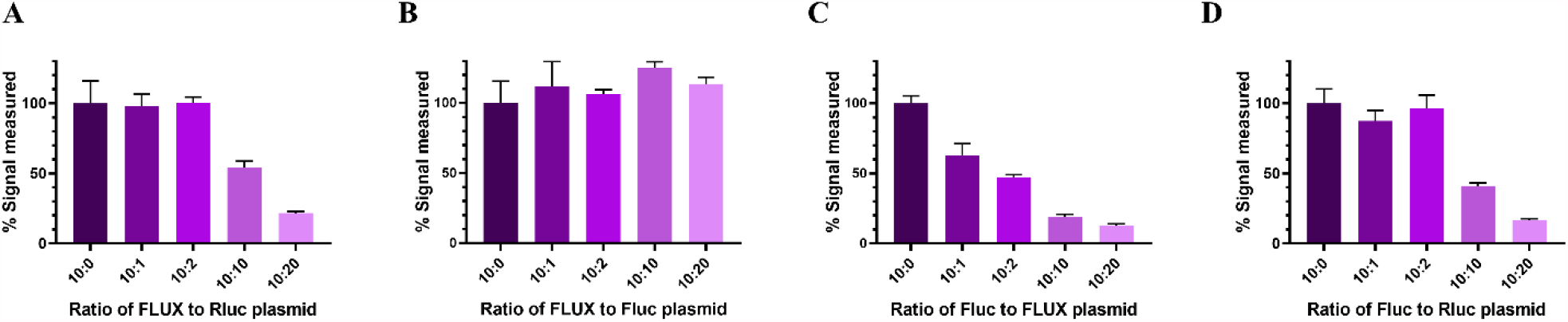
Signals of the target luciferase measured at various ratios of target:control vectors. Three types of luciferase vectors including, pGL3[luc+/SV40] vector, pGL3 [*FLUX*/SV40] and pRL-TK vectors were used as either target vector or control vector for exploring the four combination target/control vector sets including **A**. FLUX/Rluc, **B**. FLUX/Fluc, **C**. Fluc/FLUX, and **D**. Fluc/RLuc. Constant amounts of target vectors (0.07 pmol) was co-transfected into HEK293T cells with varying amounts of control vector at ratios of 10:0, 10:1, 10:2, 10:10, and 10:20. Cells were collected at 48-hours post-transfection and lysed in Passive Lysis Buffer or Lux Lysis Reagent (LLR) and luciferase activity was independently measured. The activity of FLUX was monitored by adding 100 µL of an assay reagent cocktail consisting of 5 µM FMN, 100 µM HPA, 10 µM decanal, and 100 µM NADH into a solution of cell lysate freshly mixed with 50 mU of reductase C_1_. The luminescence signal was monitored for 10 sec with a 2-sec delay using an AB-2250 single tube luminometer (ATTO Corporation, Japan). The Fluc and Rluc activities were measured using Firefly Luciferase Assay Reagent (E1500, Promega Corporation) and *Renilla* Luciferase Assay Reagent [E2810, Promega Corporation], respectively according to the manufacturer’s instructions. Percentage of signals measured (% signal measured) was calculated by dividing the signals of the target luciferase with the signals obtained in the absence of control vector (designated as 10:0 ratio) and then multiplied by 100. Error bars indicate the standard deviation (n = 4).

### Demonstrating the use of combined FLUX/Fluc reporter gene systems for investigating activators/inhibitors of the NF-κB cell signaling pathway

The nuclear factor-κB (NF-κB) transcription factor plays a critical role in inflammation, immunity, and cell proliferation, differentiation, and survival which are relevant to several diseases including cancers (Hoesel & Schmid, 2013; Oeckinghaus & Ghosh, 2009). Therefore, bioactive compounds capable of controlling signal transduction and gene regulation of this pathway are desired candidates for development of active pharmaceuticals (Oeckinghaus & Ghosh, 2009; Pahl, 1999). Luciferase reporter gene assays have been used as a valuable technique for screening NF-κB bioactive compounds because of their high sensitivity and broad range of detection. Therefore, we used the new combined FLUX/Fluc reporter system to measure the effects of Tumor Necrosis Factor (TNF) α, a known activator of the NF-κB signaling pathway, and compared it with the use of the standard Fluc/Rluc gene reporter combination.

To compare the sensitivity of both FLUX/Fluc and Fluc/Rluc reporter systems in measuring the effects of TNFα, the *FLUX* and *luc*+ reporter genes were constructed as reporter genes downstream of six tandem repeats of NF-κB transcriptional element with a TK promotor to obtain the pGL3-NF-κB[*FLUX*/TK] vector and a pGL3-NF-κB [*luc*^+^/TK] vector, respectively (Figure 6-figure supplement 1A and 1B). Combined sets of luciferase-reporter gene systems including 1. pGL3-NF-κB [*FLUX*/TK] vector with pGL3 [*luc*^*+*^/SV40] vector and 2. pGL3[*luc+*/TK] vector with pRT-TK control vector were transfected into HEK293T cells in the presence or absence of TNFα. The results showed that both sets of vector combinations gave similar signal increment responses to TNFα, 14±3-folds and 11±2-folds increment for FLUX/Fluc and Fluc/Rluc combination, respectively (Figure 6A, orange bar and purple bar, respectively). By varying concentrations of TNFα, the effects of TNFα dose-response on each reporter system can be calculated. The results showed sigmoidal dose-response curves corresponding to half-maximal response concentrations (EC50) of TNFα of 1.3±0.3ng/mL and 1.4±0.7 ng/mL for FLUX/Fluc and Fluc/Rluc combination, respectively (Figure 6B and 6C, respectively). The data clearly showed that a new combination of FLUX as the target vector and Fluc as the control vector displays similar sensitivity to the standard combined Fluc/Rluc reporter gene. It should be noted that the FLUX/Fluc gave a slightly better and consistent curve, with a lower variation value, 1.3±0.3 ng/mL *versus* 1.4±0.7 ng/mL.

**Figure 6.**
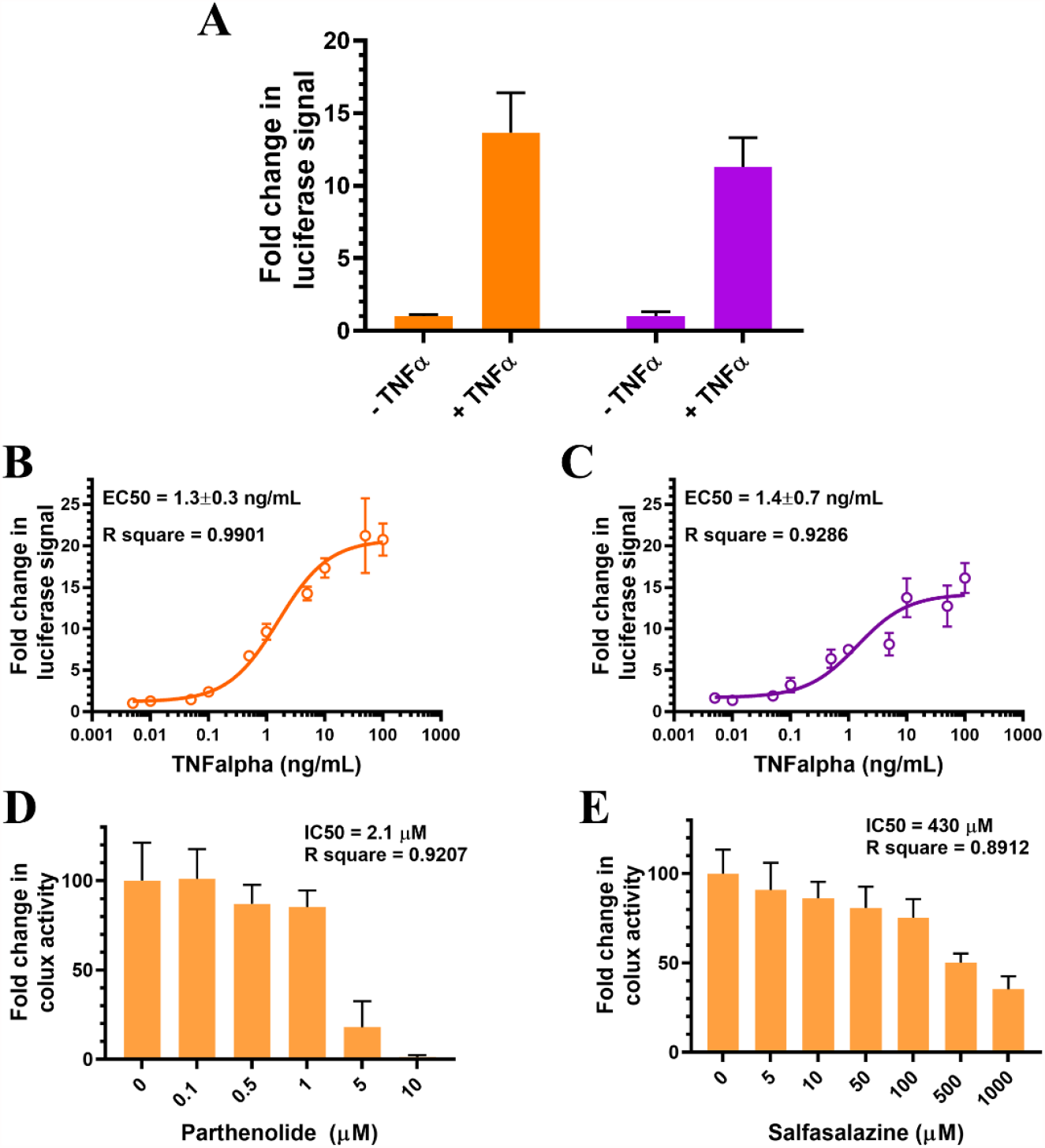
Screening of inhibitors of the NF-κB cell signaling pathway using two combined luciferase-reporter gene assays. **A** Comparison of sensitivity response of two combined luciferase-reporter gene assays including 1. FLUX/Fluc (orange) and 2 Fluc/Rluc (purple). Each combination including pGL3-NF-κB[*FLUX*/TK] vector with pGL3 [*luc*^*+*^/SV40] vector (orange) and pGL3-NF-B[*luc*^*+*^/TK] vector with pRL-TK vector (purple) were transfected into HEK293T cells. At 24 hours after transfection, the old medium was changed to fresh medium either with 10 ng/mL of TNFα (+TNFα) or without TNFα (-TNFα) for 6 hours. Cells were collected and lysed in either Passive Lysis Buffer or Lux Lysis Reagent (LLR). The luciferase activity was then measured. Error bars indicate the standard deviation (n = 4). **B and C** Investigation of TNFα dose-response of **B**. FLUX/Fluc and **C**. Fluc/Rluc combinations. Experiments were carried out by transfecting HEK293T cells that were seeded in a 6-well plate for 1-day with 1. pGL3-NFκB[*FLUX*/TK] with pGL3 [*luc*^*+*^/SV40] and 2. pGL3-NF-κB[*luc*^*+*^/TK] with pRL-TK vectors. After 12-hours of transfection, transfected cells were washed, trypsinized and seeded onto a 96-well plate. Seeded cells were further incubated for 24 hours. Then, the old medium was changed to a new medium supplied with various TNFα activators for 6 hours. Cells were collected and lysed in either Passive Lysis Buffer or Lux Lysis Reagent (LLR) and their luciferase activities were measured. **D** and **E** NF-κB inhibitor screening using the new combined FLUX/Fluc reporter gene assay. The pGL3-NF-κB[*FLUX*/TK] and pGL3 [*luc*^*+*^/SV40] vectors were transfected into HEK293T cells that were seeded in a 6-well plates for a 1-day period. After 12 hours of transfection, transfected cells were washed, trypsinized and seeded onto a 96-well plate culture plate and further incubated for 24 hours. The medium was then changed to fresh medium supplied with various concentrations of inhibitor for 30 min before activating the system by adding 10 ng/mL of TNFα for 6 hours. The cells were collected and lysed in either Passive Lysis Buffer or Lux Lysis Reagent (LLR), and their luciferase activities were measured. The activity of Lux was monitored by adding 100 µL of a reagent cocktail consisting of 5 µM FMN, 100 µM HPA, 10 µM decanal, and 100 µM NADH into cell lysate freshly mixed with 50 mU of C_1_ reductase. The luminescence signal was monitored for 10 sec with a 2-sec delay using an AB-2250 single tube luminometer (ATTO Corporation, Japan). The Fluc and Rlucactivities were measured using firefly luciferase and Renilla Luciferase Assay Reagents, E1500 and E2810, respectively [Promega Corporation] according to the manufacturer’s instructions. Error bars indicate the standard deviation (n = 4).

We further tested the ability of the new combined luciferase-reporter gene system for examining the inhibition of NF-κB using the known NF-κB inhibitors, parthenolide and sulfasalazine (Kwok et al., 2001; Weber et al., 2000). The pGL3-NF-κB [*FLUX*/TK] vector and pGL3[luc+/SV40] control vectors were transfected into HEK293T cells and the resulting transfected cells were incubated with various concentrations of the tested drugs. The results showed that half-maximum inhibitor concentrations (IC50) of parthenolide and sulfasalazine were calculated as 2.1 µM and 430 µM, respectively (Figure 6D and 6E, respectively). These values are similar to those previously reported values, 1.5 µM and 625 µM for parthenolide and sulfasalazine, respectively (Fakhrudin et al., 2014; Lakey & Cawston, 2009). Altogether, our results clearly indicate that the FLUX/Fluc reporter system can serve as an alternative luciferase-reporter gene assay which gives good precision and robustness for screening of pharmaceutical active compounds such as inhibitors of the NF-κB pathway. The low cost of FLUX (∼1/100 or ∼1/150 compared to other gene reporters) would allow researchers with limited funding to access technology without a high cost barrier, creating opportunities for life science development in poor countries.

### A general guideline for using *FLUX* as a gene reporter

A *FLUX* reporter gene consists of 2076 nucleotide bases encoding for 692 residues (Figure 1-figure supplement 1) of the optimized *lux* gene for heterologous expression in mammalian cells. The complete *FLUX* sequence is also available in the NCBI database with GenBank number MZ393808. The *FLUX* gene can be used for constructing a reporter gene in any mammalian expression vector by placing it downstream of a constitutive/inductive promotor or transcription element. For transient transfection, either the *Fluc* gene or *Rluc* gene can be used as a control for the FLUX target vector.

## DISCUSSION

This report describes the construction, optimization, validation and demonstration of applications of Lux from *V. campbellii* (*Vc*_Lux) for luciferase-reporter gene assays in mammalian cell systems. By changing the codon usage of the *Vc*_Lux gene and optimizing assay reagents, we created the <u>F</u>lavin <u>L</u>uciferase for <u>M</u>ammalian Expression (FLUX) system, which can be used as a gene reporter for mammalian cell expression. Based on detailed comparison of FLUX and the existing FLuc, FLUX shows higher signal to background (S/N) ratios than FLuc in HepG2 cells and comparable signals in other cell types. Intracellular FLUX is quite stable inside the cell for more than 72 hours, suggesting that it can be used in high throughput screening assays with high sensitivity over a broad detection range. We demonstrated that FLUX could be used as the target vector and control vector. The combination using FLUX as a target vector and Fluc as a control vector gave the most consistent signal output even better than using the combined Fluc and Rluc set. The new combined FLUX/Fluc is a sensitive detection tool which can be used for detecting Tumor Necrosis Factor (TNF)-alpha and for screening of inhibitors of the NF-κB cell signaling pathway. As the cost of FLUX system is much lower than other systems, this technology allows the use of luciferase reporter genes with a much lower price, ∼1/100 or ∼1/150 compared to other gene reporters.

The newly combined FLUX/Fluc gene-reporter system gave the most consistent signals amongst all systems tested (Figure 5B and Figure 5-figure supplement 1B). It can give consistent signals throughout a wide range of target:control ratios (10:1 up to 10:20) which is broader than the commonly used Fluc/Rluc gene reporter system (Figure 5D, and Figure 5-figure supplement 1D). The main reason behind this distinguishing feature is not clear. However, we hypothesize that this is due to stability of intracellular half-life of FLUX. The intracellular half-life of FLUX measured by inhibiting the translation process using cycloheximide was found to be much longer than 4 hours (data not shown) which is significantly longer than Fluc and RLuc (3 and 4.5 hours, respectively (Thorne et al., 2010)). Because cells have limitations for exogenous expression of all transfected genes (Hunter et al., 2019; Kaufman, 2000), high amounts of control vector for transfection may affect protein translation of the target reporter gene as is the case for data shown in Fig 5A, 5C and 5D. The long intracellular half-life of FLUX would make the system stably express its gene with less sensitivity towards the amount of the control vector used. The stable signals of FLUX after the post-transfection period (Figure 4B) also provides better advantages in transient transfection because a stable reporter gene gives less variation signal due to assay timing (Thompson et al., 1991) and is suitable as a reporter system in high throughput screening (HTP) applications because the system would be able to tolerate the long period of time needed for processing thousands of samples. The potential application of FLUX in drug screening has been shown in measurement of IC50 of the known NF-κB inhibitors, parthenolide and sulfasalazine (Figure 6D and 6E, respectively). However, the control vector of the FLUX reporter gene is not limited to only Fluc and Rluc. Researchers can employ a cheaper vector such as β-galactosidase pSV-β-Galactosidase as a control vector.

The FLUX reporter gene can be overexpressed in various types of cell-line such as Human Embryonic Kidney cells (HEK293), Mouse Embryotic Fibroblast cells (NIH3T3), Monkey Kidney Epithelial cells (COS), Human hepatocellular carcinoma cells (HepG2). The first three cell lines are the most commonly used cell lines for biomedical research (Verma et al., 2020). In particular, HEK293 cells and its derivatives have been used extensively in the transfection-based experiments, protein expression, and productions of biologics and pharmaceuticals production (Verma, 2014). They have high efficiency of transfection and protein production, and demonstrate reliable translation and processing of protein targets (Thomas & Smart, 2005). Although the ability of the FLUX reporter gene to be overexpressed in HEK293T cells was less than that in COS1, HepG2, and NIH3T3 cells, the obtained signals are sufficient for detection of dynamic changes in cell signaling in response to effectors as illustrated by measuring TNF-alpha effects on the NF-κB cell signaling pathway (Figure 5). As our experiments demonstrated that the FLUX reporter gene can function well even in the least favorable cells, the results endorse the use of FLUX system in more favorable HepG2, COS1, NIH3T3 cells. Therefore, we think that the FLUX reporter gene should be a valuable tool for transfection-based experiments and their related implications such as protein tracking, promoter screening, cell signaling and bioactive compounds screening. For example, we think that the FLUX reporter gene can be applied for detecting active compounds for other cell signaling pathways such as PI3K/Akt and JAK/STAT signaling pathways that regulate cell growth, proliferation, migration, and apoptosis, which are critical process for tumorigenesis that is a serious problems in low- to middle-income countries (Gelband et al., 2015; Luo et al., 2003; Rawlings et al., 2004). Moreover, our results suggest that FLUX can be used in replacement of Fluc in various applications such as in the systems where compounds such as resveratrol and benzothiazoles derivatives, known inhibitors of Fluc, are present (Bakhtiarova et al., 2006; Braeuning & Vetter, 2012; Leitão & da Silva, 2010).

We also noted that the FLUX reporter gene gives outstanding signals in HepG2 cells, yielding much higher S/N than that the Fluc reporter gene (Figure 3). HepG2 is a human hepatocellular carcinoma cell which has drug metabolizing enzymes comparable to normal hepatocytes. The cells can be cultivated *in vitro* and used for drug testing conveniently (Castell et al., 2006). Although HepG2 is less used for investigating cellular signaling and events than other cell types, it is commonly used for drug cytotoxicity screening and investigating drug toxic mechanisms. We hope that the FLUX reporter gene developed here which can be expressed well in HepG2 cells can contribute to future drug cytotoxicity screening and many other applications related to this cell-line.

## MATERIALS AND METHODS

### Key Resources Table

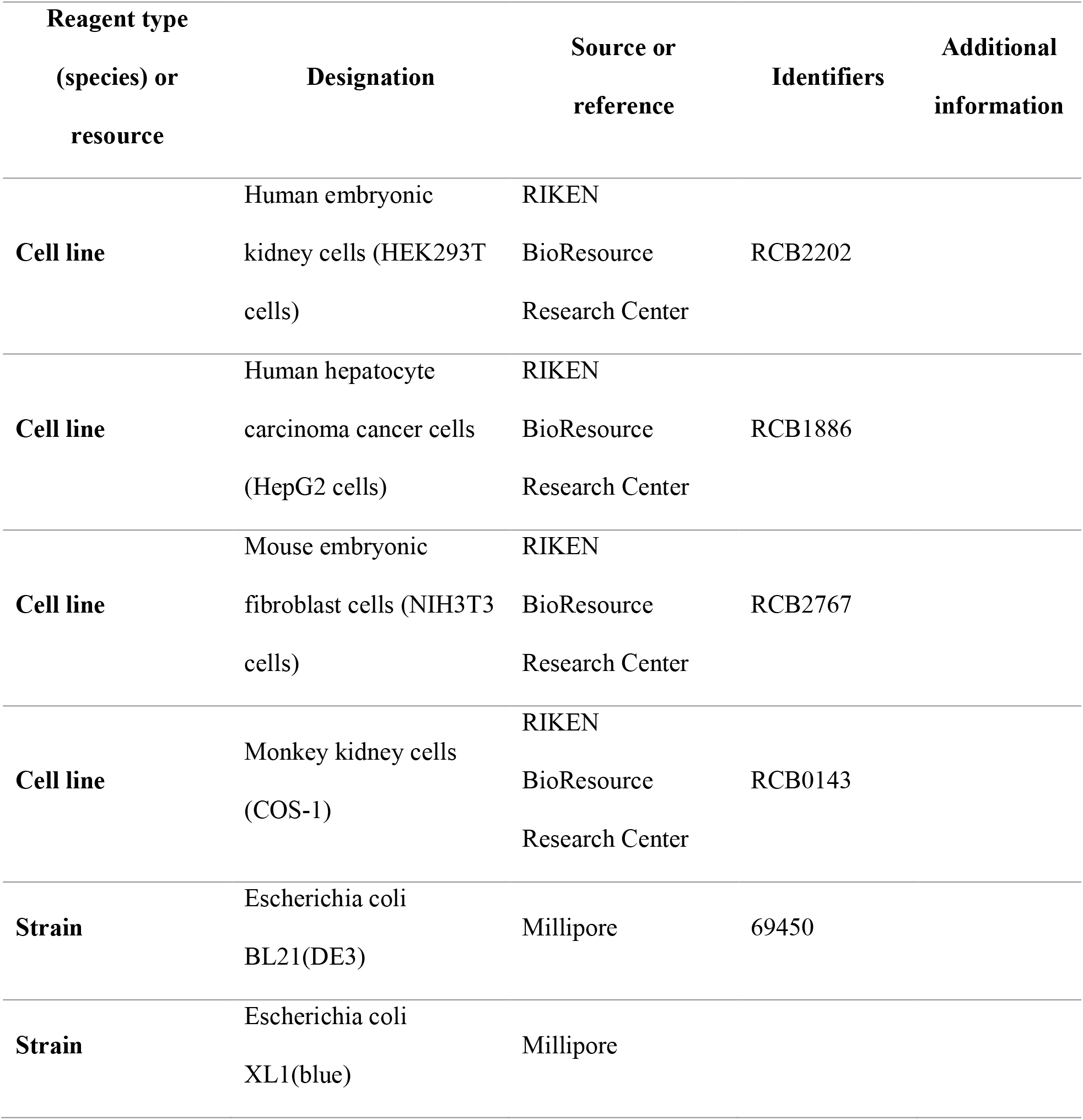
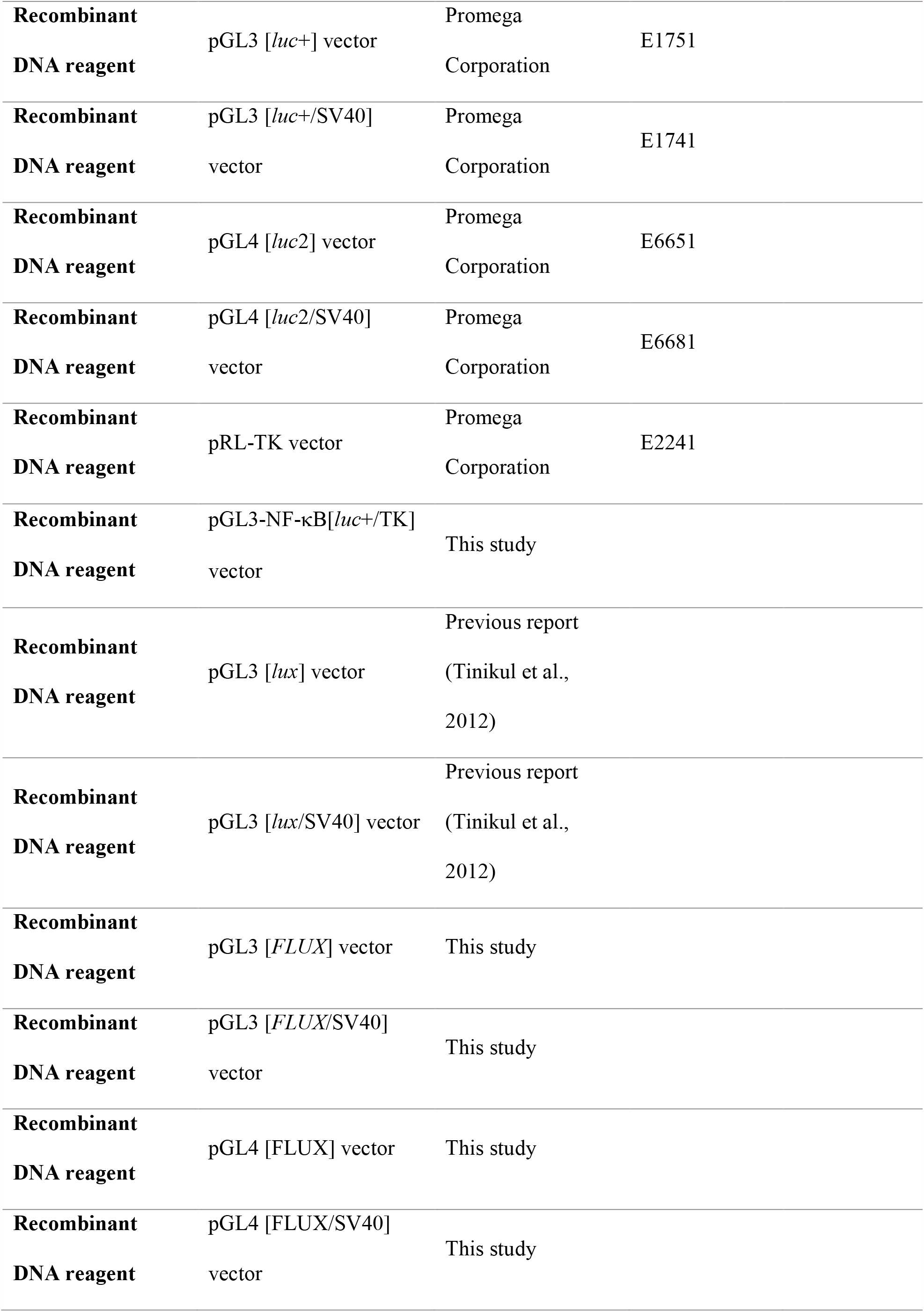
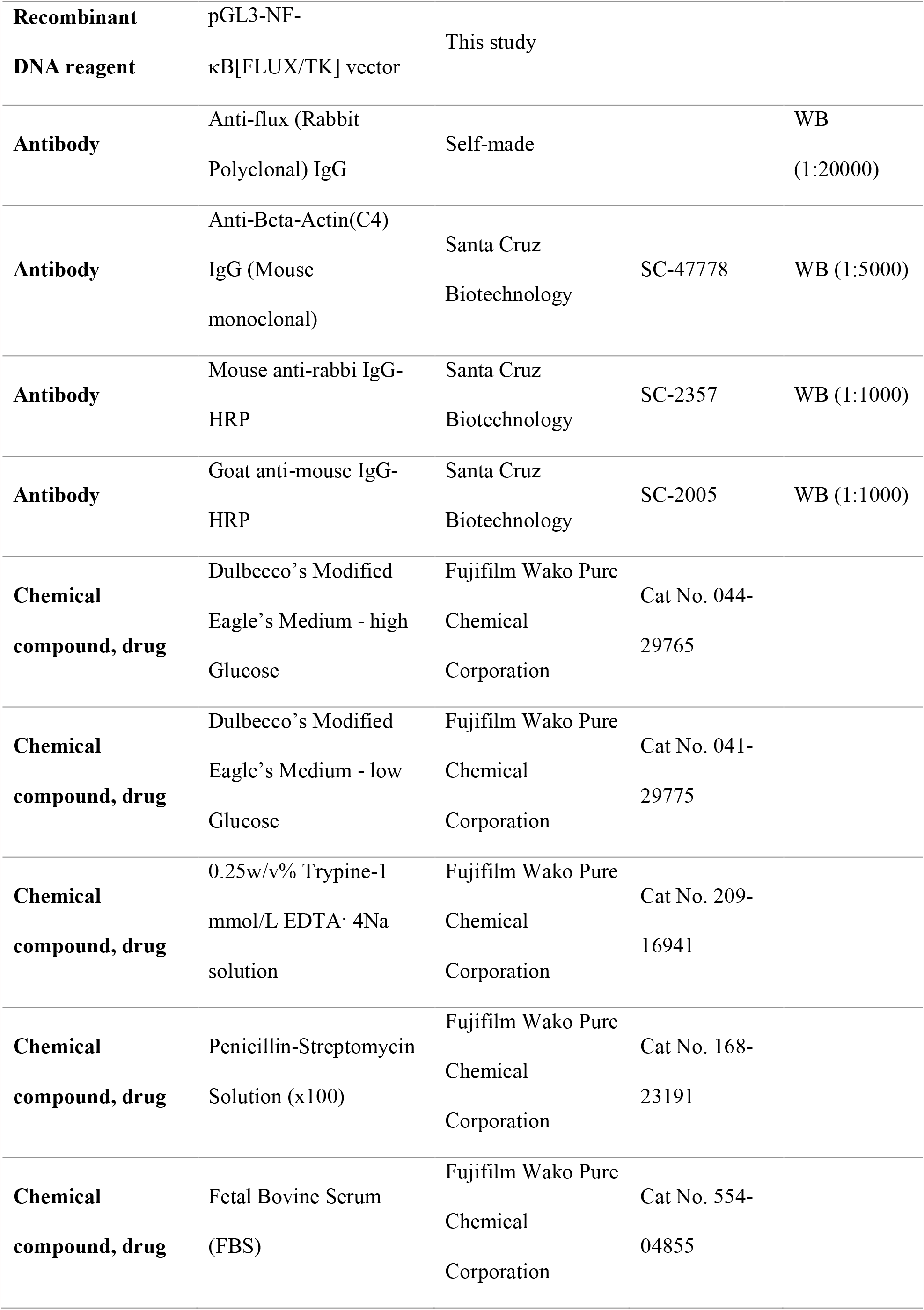
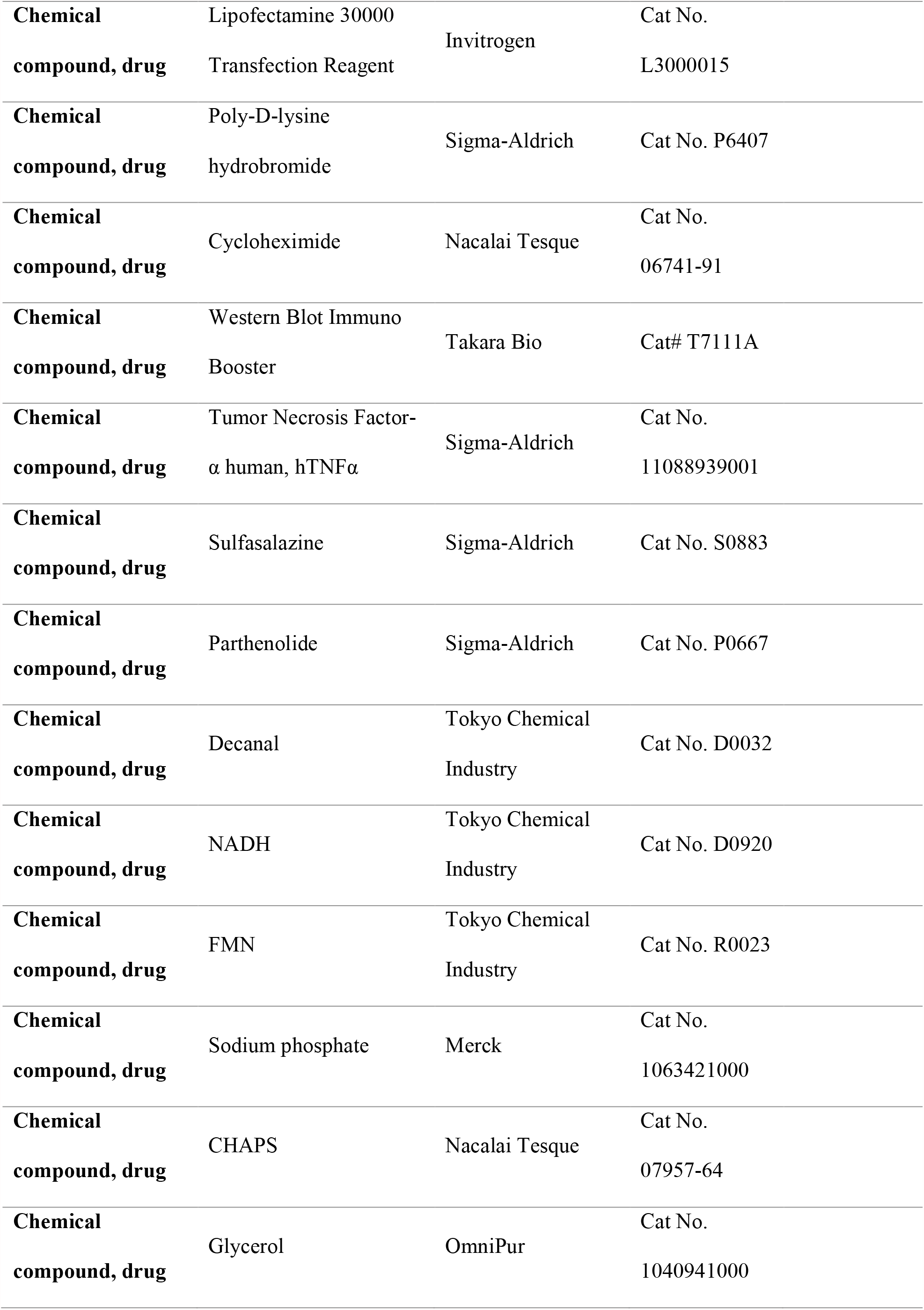
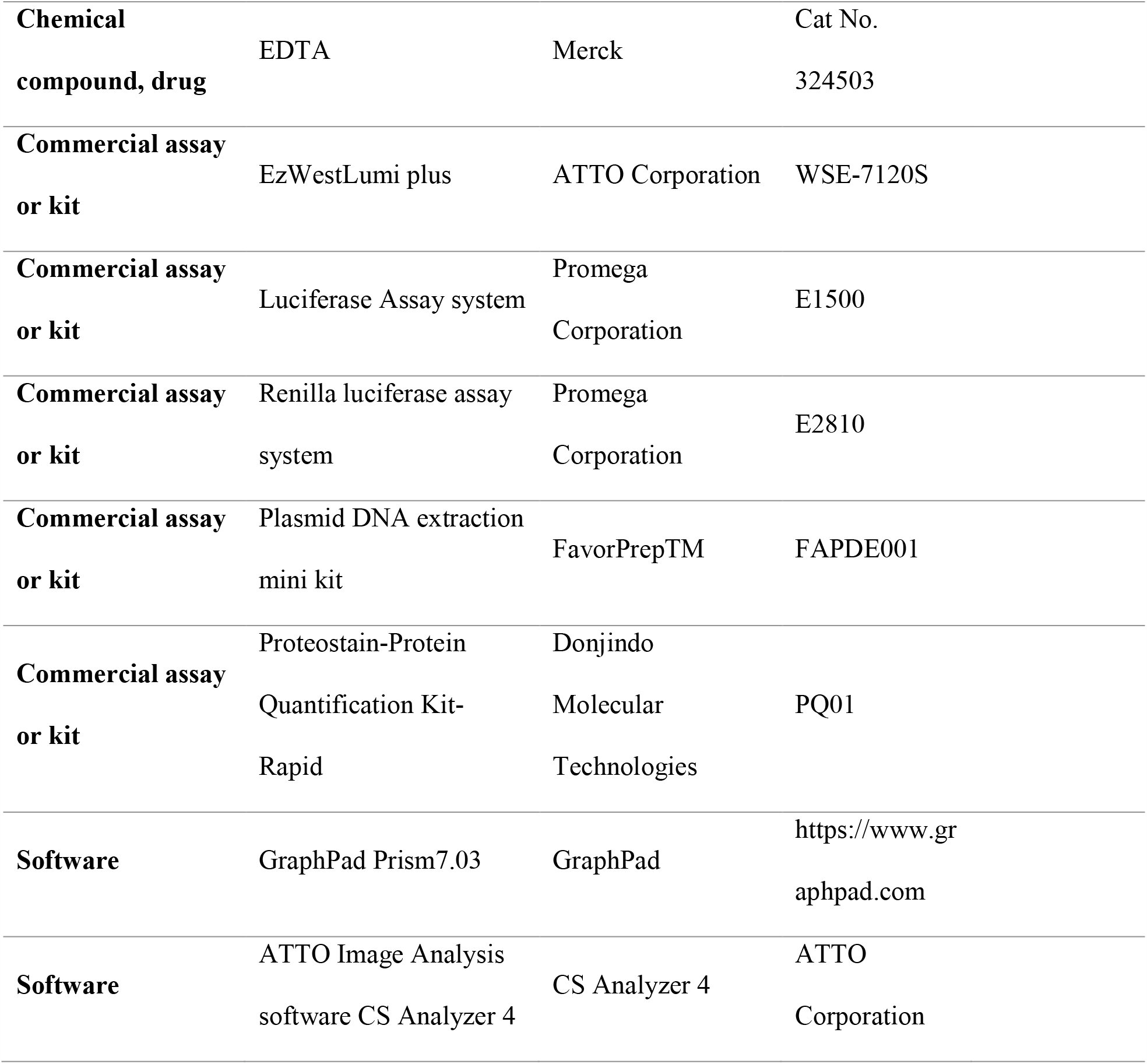

| Reagent type<br>(species) or<br>resource | Designation | Source or<br>reference | Identifiers | Additional<br>information |
| --- | --- | --- | --- | --- |
| Cell line | Human embryonic | RIKEN |  |  |
|  | kidney cells (HEK293T | BioResource | RCB2202 |  |
|  | cells) | Research Center |  |  |
| Cell line | Human hepatocyte | RIKEN |  |  |
|  | carcinoma cancer cells | BioResource | RCB1886 |  |
|  | (HepG2 cells) | Research Center |  |  |
| Cell line | Mouse embryonic | RIKEN |  |  |
|  | fibroblast cells (NIH3T3 | BioResource | RCB2767 |  |
|  | cells) | Research Center |  |  |
| Cell line | Monkey kidney cells<br>(COS-1) | RIKEN |  |  |
|  |  | BioResource | RCB0143 |  |
|  |  | Research Center |  |  |
| Strain | Escherichia coli<br>BL21(DE3) | Millipore | 69450 |  |
| Strain | Escherichia coli<br>XL1(blue) | Millipore |  |  |

|  |  |  |  |
| --- | --- | --- | --- |
| **Recombinant**<br>**DNA reagent** | pGL3 [*luc*+] vector | Promega<br>Corporation | E1751 |
| **Recombinant**<br>**DNA reagent** | pGL3 [*luc*+/*SV40*]<br>vector | Promega<br>Corporation | E1741 |
| **Recombinant**<br>**DNA reagent** | pGL4 [*luc2*] vector | Promega<br>Corporation | E6651 |
| **Recombinant**<br>**DNA reagent** | pGL4 [*luc2*/*SV40*]<br>vector | Promega<br>Corporation | E6681 |
| **Recombinant**<br>**DNA reagent** | pRL-TK vector | Promega<br>Corporation | E2241 |
| **Recombinant**<br>**DNA reagent** | pGL3-NF-κB[*luc*+/*TK*]<br>vector | This study | |
| **Recombinant**<br>**DNA reagent** | pGL3 [*lux*] vector | Previous report<br>(Tinikul et al.,<br>2012) |  |
| **Recombinant**<br>**DNA reagent** | pGL3 [*lux*/*SV40*] vector | Previous report<br>(Tinikul et al.,<br>2012) |  |
| **Recombinant**<br>**DNA reagent** | pGL3 [*FLUX*] vector | This study |  |
| **Recombinant**<br>**DNA reagent** | pGL3 [*FLUX*/*SV40*]<br>vector | This study |  |
| **Recombinant**<br>**DNA reagent** | pGL4 [*FLUX*] vector | This study |  |
| **Recombinant**<br>**DNA reagent** | pGL4 [*FLUX*/*SV40*]<br>vector | This study |  |
| **Recombinant DNA reagent** | pGL3-NF-κB[FLUX/TK] vector | This study |  |  |
| **Antibody** | Anti-flux (Rabbit Polyclonal) IgG | Self-made |  | WB<br>(1:20000) |
| **Antibody** | Anti-Beta-Actin(C4) IgG (Mouse monoclonal) | Santa Cruz Biotechnology | SC-47778 | WB (1:5000) |
| **Antibody** | Mouse anti-rabbi IgG-HRP | Santa Cruz Biotechnology | SC-2357 | WB (1:1000) |
| **Antibody** | Goat anti-mouse IgG-HRP | Santa Cruz Biotechnology | SC-2005 | WB (1:1000) |
| **Chemical compound, drug** | Dulbecco's Modified Eagle's Medium - high Glucose | Fujifilm Wako Pure Chemical Corporation | Cat No. 044-29765 |  |
| **Chemical compound, drug** | Dulbecco's Modified Eagle's Medium - low Glucose | Fujifilm Wako Pure Chemical Corporation | Cat No. 041-29775 |  |
| **Chemical compound, drug** | 0.25w/v% Trypine-1 mmol/L EDTA· 4Na solution | Fujifilm Wako Pure Chemical Corporation | Cat No. 209-16941 |  |
| **Chemical compound, drug** | Penicillin-Streptomycin Solution (x100) | Fujifilm Wako Pure Chemical Corporation | Cat No. 168-23191 |  |
| **Chemical compound, drug** | Fetal Bovine Serum (FBS) | Fujifilm Wako Pure Chemical Corporation | Cat No. 554-04855 |  |

|  |  |  |  |
| --- | --- | --- | --- |
| **Chemical**<br>**compound, drug** | Lipofectamine 30000<br>Transfection Reagent | Invitrogen | Cat No.<br>L3000015 |
| **Chemical**<br>**compound, drug** | Poly-D-lysine<br>hydrobromide | Sigma-Aldrich | Cat No. P6407 |
| **Chemical**<br>**compound, drug** | Cycloheximide | Nacalai Tesque | Cat No.<br>06741-91 |
| **Chemical**<br>**compound, drug** | Western Blot Immuno<br>Booster | Takara Bio | Cat# T7111A |
| **Chemical**<br>**compound, drug** | Tumor Necrosis Factor-<br>$\alpha$ human, hTNFα | Sigma-Aldrich | Cat No.<br>11088939001 |
| **Chemical**<br>**compound, drug** | Sulfasalazine | Sigma-Aldrich | Cat No. S0883 |
| **Chemical**<br>**compound, drug** | Parthenolide | Sigma-Aldrich | Cat No. P0667 |
| **Chemical**<br>**compound, drug** | Decanal | Tokyo Chemical<br>Industry | Cat No. D0032 |
| **Chemical**<br>**compound, drug** | NADH | Tokyo Chemical<br>Industry | Cat No. D0920 |
| **Chemical**<br>**compound, drug** | FMN | Tokyo Chemical<br>Industry | Cat No. R0023 |
| **Chemical**<br>**compound, drug** | Sodium phosphate | Merck | Cat No.<br>1063421000 |
| **Chemical**<br>**compound, drug** | CHAPS | Nacalai Tesque | Cat No.<br>07957-64 |
| **Chemical**<br>**compound, drug** | Glycerol | OmniPur | Cat No.<br>1040941000 |
| **Chemical compound, drug** | EDTA | Merck | Cat No.<br>324503 |
| **Commercial assay or kit** | EzWestLumi plus | ATTO Corporation | WSE-7120S |
| **Commercial assay or kit** | Luciferase Assay system | Promega Corporation | E1500 |
| **Commercial assay or kit** | Renilla luciferase assay system | Promega Corporation | E2810 |
| **Commercial assay or kit** | Plasmid DNA extraction mini kit | FavorPrep™ | FAPDE001 |
| **Commercial assay or kit** | Proteostain-Protein Quantification Kit-Rapid | Donjindo Molecular Technologies | PQ01 |
| **Software** | GraphPad Prism7.03 | GraphPad | https://www.graphpad.com |
| **Software** | ATTO Image Analysis software CS Analyzer 4 | CS Analyzer 4 | ATTO Corporation |

### Mammalian cell culture, transient transfection, and cell harvesting

Human Embryotic Kidney (HEK293T) [RCB2202] cells and human hepatocyte carcinoma cancer cells (HepG2 cells) [RCB1886] were grown in Dulbecco’s Modified Eagle Medium (DMEM)-low glucose supplemented with 10% (v/v) heat-inactivated fetal bovine serum plus 1% (w/v) penicillin-streptomycin. Mouse embryonic fibroblast cells (NIH3T3 cells) [RCB2767] and monkey kidney cells (COS1) [RCB0143] were grown in DEME-high glucose supplemented with 10% (v/v) heat-inactivated fetal bovine serum plus 1% (w/v) penicillin-streptomycin. Four types of cells were maintained at 37 °C with 5% CO_2_. For transient transfection, 1×10^5^ cells per well were plated and cultured in suitable DEME media supplemented with 10% (v/v) heat-inactivated fetal bovine serum in 24 well plates for one day prior to transfection. Cells were transfected by adding 0.07 pmol of target vector and 0.007 pmol of for control vector with lipofectamine^™^3000 in DEME-free serum. The transfected cells were maintained for 24-72 hours. Cells were harvested by washing cells with 500 µl PBS buffer pH 7.4 and detached cells were collected using specific lysis buffer. A 1x Passive Lysis Buffer was used for collecting Fluc transfected cell while a 1x Lux Lysis Reagent (LLR) was used for collecting FLUX transfected cells. The lysate was kept at -80 °C until used.

### Protein extraction and protein concentration determination

Cells were lysed by a freezing-thawing process (freezing at -80 °C for 10 min and thawing at room temperature water for 2 min). The cell lysate was collected by centrifugation at 12000 rpm at 4 °C for 10 min. Total protein concentration was determined using Proteostain-Protein Quantification Rapid-Kit (Donjindo Molecular Technologies) according to the manufacturer’s instructions. The absorbance change at 595 nm was measured using a microplate reader spectrophotometer (iMark^™^, BioRad). A plot of absorbance change *versus* BSA concentration was used as a standard curve. Protein concentrations of cell lysate were calculated based on the standard curve.

### Measurement of luciferase activity in cell lysate

Activity of luciferase was assayed by monitoring light emission using AB-2250 single tube luminometer (ATTO Corporation, Japan). The Lux cocktail reagent consisting of 5 µM FMN, 10 µM decanal, 100 µM *p*-hydroxyphenylacetic acid (HPA), and 100 µM NADH in 50 mM sodium phosphate buffer pH 7.0. was injected into a reaction chamber containing 2-10 µL of cell lysate and 50 mU of C_1_ reductase. The light emission was monitored for 10 second with 2 second delay. Fluc and Rluc activities were measured using firefly luciferase and Renilla Luciferase Assay Reagents, E1500 and E2810 (Promega Corporation, Wisconsin, USA), respectively according to the manufacturer’s instructions. The integrated peak area was reported as relative light units (RLU). Signal of luciferase-based experiments were normalized using light emitted from the control vector.

### Measurement of purified Lux activity

Activity of purified Lux was assayed by monitoring light emission using an AB-2250 single tube luminometer (ATTO Corporation, Japan). A Lux cocktail reagent consisting of 5 µM FMN, 10 µM decanal, 100 µM *p*-hydroxyphenylacetic acid (HPA), and 100 µM NADH in in 50 mM sodium phosphate buffer pH 7.0. was injected into a reaction chamber consisting of 2 µL of purified Lux and 50 mU of C_1_ reductase. The light emission was monitored for 10 second with 2 second delay. The integrated peak area was reported as relative light units (RLU).

### Western blot analysis

The cell lysate was separated by 12.5% (w/v) SDS-PAGE electrophoresis (ATTO Cooperation, Japan) and transferred onto a PVDF membrane (Bio-rad, USA). The membrane blots were blocked in 1x EzBlockChemi (ATTO Cooperation, Japan) for 1 hour at room temperature and then incubated with primary antibody (Anti-flux IgG / Anti-β-actin IgG) which was diluted in Solution I of Western BLoT Immuno Booster (ATTO Cooperation, Japan) for 24 hours. at 4 °C. The membrane was washed by TBS-T for 3 times before incubating in HRP-conjugated secondary antibodies (mouse-anti-rabbit HRP / goat-anti-mouse HRP) for 1 hour at room temperature. The membrane was washed 3-times using TBS-T before incubating in chemiluminescent reagent for HRP (EzWestLumi plus, ATTO Cooperation, Japan). The chemiluminescence was detected by WSE-6100 LuminoGraph I Gel documentation system (ATTO Cooperation, Japan). The band intensity was measured by ATTO Image Analysis software CS Analyzer 4 (ATTO Cooperation, Japan).

### Validation of novel dual-luciferase assay using NF-κB transcription element

Four sets of validated reporter genes (Table 1) were independently transfected into 5×10^5^ HEK293T cells that were plated one day prior to transfection in 6-well plates using lipofectamine^™^3000 in DEME-free serum. After 12 hours post-transfection, the transfected cells were washed, trypsinized, and seeded on a 96-well plate. The culture plate was incubated for 24 hours at 37 °C with 5% CO_2_. The culture medium was then changed to a new medium either supplied with 0.005-10 ng/mL of TNFα or without TNFα. The transfected cells were continuously stimulated for 6 hours. Cells were harvested by washing cells with 500 µl PBS buffer pH 7.4 and collecting using specific lysis regent. A 20-50 µL of 1x Passive Lysis Buffer was used for collecting firefly luciferase transfected cell while 20-50 µL of 1x Lux Lysis Reagent (LLR) was used for collecting bacterial luciferase transfected cells. The luciferase activity was independently measured according to the measurement protocol described above.

**Table 1.** Four combinations of reporter genes for validating the function of FLUX reporter gene

| Condition | NF-κB reporter vector | Internal control vector |
| --- | --- | --- |
| 1 | pGL3-NF-κB[*luc*<sup>+</sup>/TK] | pRL-TK vector |
| 2 | pGL3-NF-κB[*FLUX*/TK] | pRL-TK vector |
| 3 | pGL3-NF-κB[*luc*<sup>+</sup>/TK] | pGL3 [*FLUX*/SV40] |
| 4 | pGL3-NF-κB[*FLUX*/TK] | pGL3[*luc*<sup>+</sup>/SV40] |

### Use of a new combined FLUX/Fluc for demonstrating investigation of inhibitors of the NF-κB signaling pathway

The pGL3-NF-κB[*FLUX*/TK] and pGL3 [*luc+*/SV40] vectors were co-transfected into 5×10^5^ HEK293T cells that were plated one day prior to transfection in 6-well plates using lipofectamine^™^3000 in DEME-free serum. After 12 hours post-transfection, the transfected cells were washed, trypsinized, and seeded on a 96-well plate. The culture plate was incubated for 24 hours at 37 °C with 5% CO_2_. The culture medium was then changed to a new medium supplied with various concentrations of inhibitors for 30 min before stimulating the system by adding 10 ng/mL of TNFα for 6 hours. The cells were then collected by adding 50 µL of Lux Lysis Reagent (LLR) and luciferase activities were independently measured. The activity of Lux was monitored by adding 100 µL of a cocktail reagent consisting of 5 µM FMN, 100 µM HPA, 10 µM decanal, and 100 µM NADH into 2-10 µL of cell lysate freshly mixed with 50 mU of C_1_ reductase. The luminescence signal was monitored for 10 sec with a 2 sec delay using an AB-2250 single luminometer (ATTO Corporation, Japan). The Fluc activity was measured using firefly Luciferase Assay Reagent [E2810, Promega Corporation, USA] according to the manufacturer’s instructions.

## ACKNOWLEDGMENTS

This work was supported by grants from Vidyasrimedhi Institute of Science and Technology (VISTEC) and from the Global Partnership grant from Program Management Unit-B (to P.C. and J. P.) and partially supported by the National Institute of Advanced Industrial Science and Technology (AIST) to Y.O., and the Central Instrument Facility (CIF), Faculty of Science, Mahidol University to R.T. We would like to thank Frank Fan, Promega company for the generous gift of pGL4.13[luc2/SV40] vector and pGL4.10[luc2] vectors.

