## Supplemental Figures for "High Sensitivity and Low-Cost Flavin luciferase (FLUX)-based Reporter Gene for Mammalian Cell Expression"

### Figure supplement

1 ATGAAGTTTGGCAATTTCCTGCTGACATACCAGCCTCCTGAACTGAGCCAGACCGAAGTG 60
1 M K F G N F L L T Y Q P P E L S Q T E V 20
61 ATGAAGAGACTGGTGAACCTGGGCAAGGCCTCTGAGGGGTGCGGATTCGACACAGTGTGG 120
21 M K R L V N L G K A S E G C G F D T V W 40
121 CTGCTGGAACACCACTTCACCGAGTTCGGACTGCTGGGAAACCCATACGTGGCAGCTGCA 180
41 L L E H H F T E F G L L G N P Y V A A A 60
181 CACCTGCTGGGAGCAACTGAAAAGCTGAATGTGGGAACCGCCGCTATCGTCCTGCCCACA 240
61 H L L G A T E K L N V G T A A I V L P T 80
241 GCTCATCCTGTGAGGCAGGCAGAGGACGTGAACCTGCTGGATCAGATGTCAAAAGGCAGG 300
81 A H P V R Q A E D V N L L D Q M S K G R 100
301 TTCCGCTTTGGGATTTGCCGCGGACTGTATGACAAGGATTTTCGAGTGTTCGGGACCGAC 360
101 F R F G I C R G L Y D K D F R V F G T D 120
361 ATGGATAATAGCCGGGCCCTGATGGACTGTTGGTACGATCTGATGAAGGAAGGCTTTAAC 420
121 M D N S R A L M D C W Y D L M K E G F N 140
421 GAGGGGTATATCGCAGCCGACAATGAGCACATCAAGTTCCCTAAAATTCAGCTGAACCCA 480
141 E G Y I A A D N E H I K F P K I Q L N P 160
481 TCCGCCTACACACAGGGAGGGGCTCCAGTGTATGTGGTCGCCGAATCTGCTAGTACCACA 540
161 S A Y T Q G G A P V Y V V A E S A S T T 180
541 GAATGGGCTGCAGAGAGAGGACTGCCCATGATCCTGTCTTGGATCATTAACACACATGAG 600
181 E W A A E R G L P M I L S W I I N T H E 200
601 AAGAAAGCCCAGCTGGATCTGTACAATGAAGTGGCTACTGAGCACGGCTATGACGTGACC 660
201 K K A Q L D L Y N E V A T E H G Y D V T 220
661 AAAATCGATCATTGCCTGTCCTACATTACTTCTGTGGACCACGATAGCAACAAGGCTAAA 720
221 K I D H C L S Y I T S V D H D S N K A K 240
721 GACATCTGTAGGAATTTTCTGGGGCATTGGTACGATAGTTATGTGAACGCCACCAAGATT 780
241 D I C R N F L G H W Y D S Y V N A T K I 260
781 TTCGACGATTCAGACCAGACAAAGGGATATGACTTCAACAAGGGCCAGTGGCGCGACTTC 840
261 F D D S D Q T K G Y D F N K G Q W R D F 280
841 GTGCTGAAGGGCCACAAAGACACTAACCGGAGAATCGATTACAGCTATGAGATTAATCCA 900
281 V L K G H K D T N R R I D Y S Y E I N P 300
901 GTGGGAACCCCCGAGGAATGTATCGCTATCATTCAGCAGGACATTGATGCAACAGGCATC 960
301 V G T P E E C I A I I Q Q D I D A T G I 320

```
961 AACAATATTTGCTGTGGGTTTGAAGCAAACGGCAGCGAGGAAGAGATCATTGCCTCTATG 1020
321  N  N  I  C  C  G  F  E  A  N  G  S  E  E  E  I  I  A  S  M  340
1021 AAGCTGTTCCAGAGTGACGTGATGCCTTACCTGAAGGAGAAACAGGTCATCAATATTTTT 1080
341  K  L  F  Q  S  D  V  M  P  Y  L  K  E  K  Q  V  I  N  I  F  360
1081 GAAAAGGAGAGGGATCAGAAATTTGGCCTGTTCTTTCTGAACTTCATGAATAGTAAACGC 1140
361  E  K  E  R  D  Q  K  F  G  L  F  F  L  N  F  M  N  S  K  R  380
1141 AGCTCCGACCAGATCATTGAAGAGATGCTGGATACAGCTCACTACGTGGACCAGCTGAAG 1200
381  S  S  D  Q  I  I  E  E  M  L  D  T  A  H  Y  V  D  Q  L  K  400
1201 TTTGATACTCTGGCCGTCTATGAGAACCATTTCTCCAACAATGGAGTGGTCGGCGCCCCA 1260
401  F  D  T  L  A  V  Y  E  N  H  F  S  N  N  G  V  V  G  A  P  420
1261 CTGACTGTGGCTGGATTCCTGCTGGGCATGACCAAGAACGCAAAAGTGGCCTCTCTGAAT 1320
421  L  T  V  A  G  F  L  L  G  M  T  K  N  A  K  V  A  S  L  N  440
1321 CACGTCATCACTACCCACCATCCCGTGCGGGTCGCTGAAGAGGCATGCCTGCTGGACCAG 1380
441  H  V  I  T  T  H  H  P  V  R  V  A  E  E  A  C  L  L  D  Q  460
1381 ATGAGTGAAGGCAGATTCGTGTTTGGGTTCAGTGACTGTGAGAAGTCAGCCGATATGCGA 1440
461  M  S  E  G  R  F  V  F  G  F  S  D  C  E  K  S  A  D  M  R  480
1441 TTCTTTAACCGGCCCACCGATTCACAGTTTCAGCTGTTCAGCGAGTGCCACAAAATCATT 1500
481  F  F  N  R  P  T  D  S  Q  F  Q  L  F  S  E  C  H  K  I  I  500
1501 AATGACGCCTTTACAACTGGCTACTGTCATCCTAACAATGACTTCTACAGCTTCCCCAAG 1560
501  N  D  A  F  T  T  G  Y  C  H  P  N  N  D  F  Y  S  F  P  K  520
1561 ATCTCCGTGAACCCTCACGCCTTTACCGAGGGAGGCCCTGCACAGTTCGTCAATGCCACA 1620
521  I  S  V  N  P  H  A  F  T  E  G  G  P  A  Q  F  V  N  A  T  540
1621 AGCAAGGAAGTGGTCGAGTGGGCCGCTAAACTGGGCCTGCCACTGGTGTTCAAGTGGGAC 1680
541  S  K  E  V  V  E  W  A  A  K  L  G  L  P  L  V  F  K  W  D  560
1681 GATTCCAATGCACAGCGAAAAGAATACGCCGGCCTGTATCACGAGGTGGCACAGGCCCAC 1740
561  D  S  N  A  Q  R  K  E  Y  A  G  L  Y  H  E  V  A  Q  A  H  580
1741 GGCGTGGACGTGAGCCAGGTCCGCCATAAGCTGACTCTGCTGGTGAACCAGAATGTCGAT 1800
581  G  V  D  V  S  Q  V  R  H  K  L  T  L  L  V  N  Q  N  V  D  600
1801 GGCGAAGCAGCCAGAGCTGAGGCAAGGGTGTACCTGGAAGAGTTTGTCCGGGAAAGCTAT 1860
601  G  E  A  A  R  A  E  A  R  V  Y  L  E  E  F  V  R  E  S  Y  620
1861 CCAAACACCGACTTCGAGCAGAAAATGGTGGAACTGCTGTCCGAGAATGCTATCGGGACC 1920
621  P  N  T  D  F  E  Q  K  M  V  E  L  L  S  E  N  A  I  G  T  640
1921 TACGAAGAGTCTACACAGGCTGCAAGAGTGGCCATTGAGTGCTGTGGAGCCGCTGACCTG 1980
641  Y  E  E  S  T  Q  A  A  R  V  A  I  E  C  C  G  A  A  D  L  660
```

```
1981 CTGATGTCATTCGAAAGCATGGAGGATAAGGCACAGCAGAGAGCCGTGATTGATGTCGTG 2040
 661  L  M  S  F  E  S  M  E  D  K  A  Q  Q  R  A  V  I  D  V  V  680
2041 AACGCAAATATCGTGAAATACCACTCATAA 2070
 681  N  A  N  I  V  K  Y  H  S  *  690
```

**Figure 1-figure supplement 1.** Nucleotide (2070 base pairs) and deduced amino acid (690 residues) sequences of the FLUX gene. Highlighted in yellow is an artificial peptide linker which joins the *luxA* and *luxB* genes.

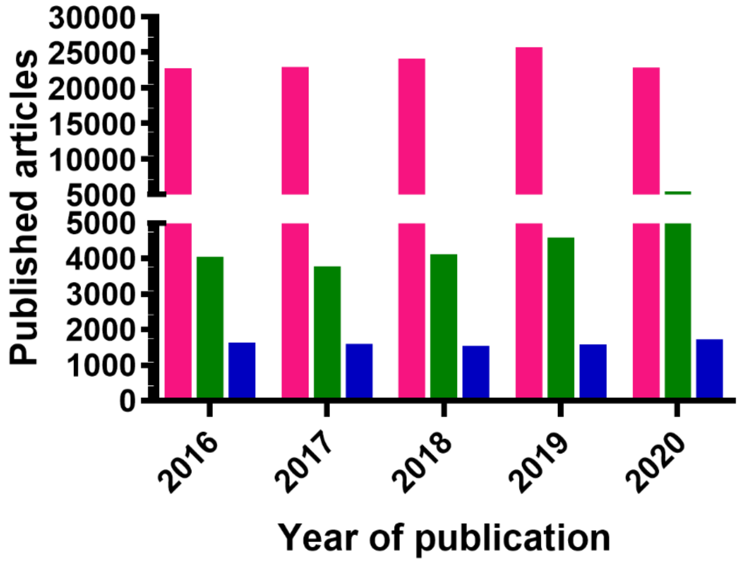


**Figure 3-figure supplement 1.** Published articles using all types of luciferase reporters (pink bar), the pGL3 vector (green bar) and the pGL4 vector (blue bar). Keywords containing either luciferase reporter, pGL3 or pGL4 were searched in google scholar to find related articles during 2016-2020.

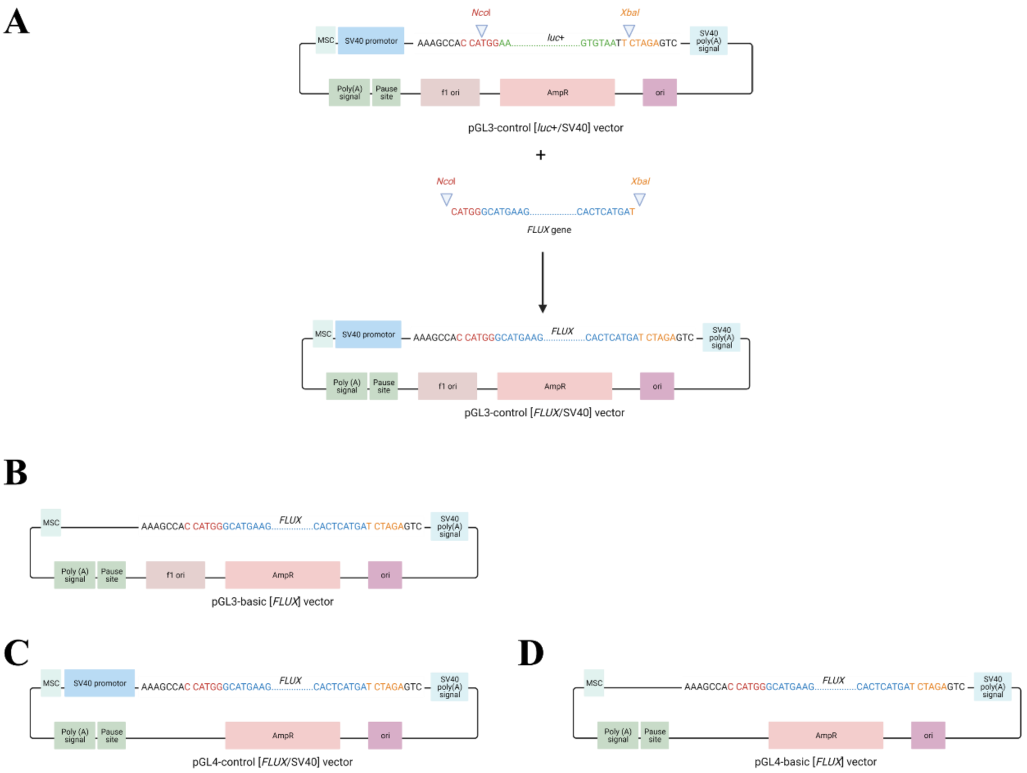


**Figure 3-figure supplement 2.** Maps of FLUX genes constructed in two types of pGL vectors in the presence and absence of a constitutive SV40 promoter. **A.** The pGL3 [*FLUX*/SV40] vector **B.** The pGL3 [*FLUX*] vector **C.** The pGL4 [*FLUX*/SV40] vector and **D.** The pGL4 [*FLUX*] vector. The construction processes of each vector were all similar except for the pGL4 [*FLUX*/SV40] vector. First, the pGL vector containing the Fluc gene was digested with *Nco*I and *Xba*I restriction enzymes to remove the *Fluc* gene. The *FLUX* gene was amplified and digested by *Nco*I and *Xba*I restriction enzymes. The digested pGL vector was then ligated with the digested *FLUX* gene to yield the pGL vector consisting of FLUX as a reporter gene.

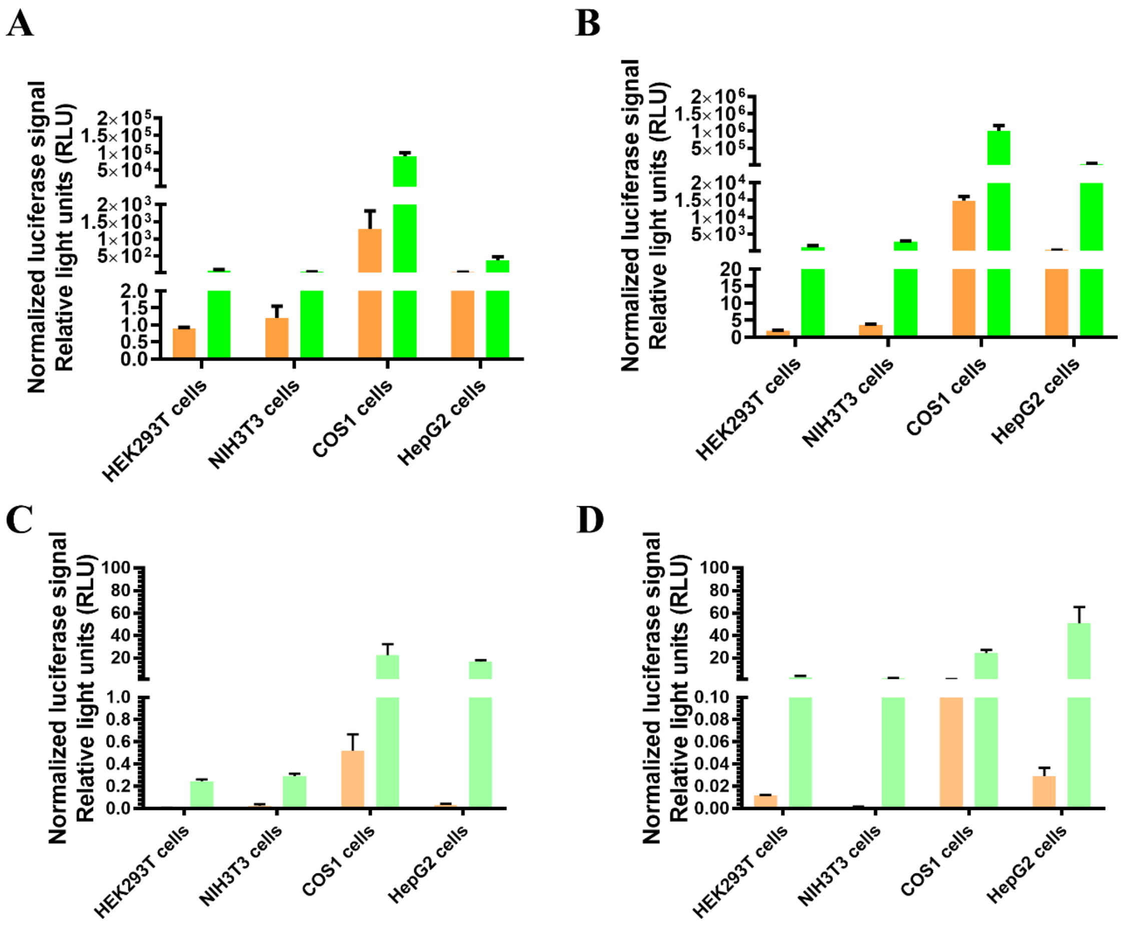


**Figure 3-figure supplement 3.** Comparison of normalized luciferase signals between FLUX (orange bar) and Fluc (green bar) in various cell types including HEK293T, NIH3T3, COS1 and HepG2 cells. **A.** pGL3 backbone vectors in the presence or **C.** absence of a constitutive SV40 promotor. **B.** pGL4 backbone vectors in the presence or **D.** absence of a constitutive SV40 promotor. Each reporter gene was co-transfected with the pRL-TK vector as an internal control for each cell type. Cells were collected at 48 hours post-transfection in either Passive Lysis Buffer or Lux Lysis Reagent (LLR) and luciferase activities were measured. The activity of Lux was monitored by adding 100 µL of a cocktail reagent consisting of 5 µM FMN, 100 µM HPA, 10 µM decanal and 100 µM NADH into a cell lysate which was freshly mixed with 50 mU of $C_1$ reductase. The luminescence signal was monitored for 10 sec with a 2-sec delay using an AB-2250 single tube luminometer (ATTO Corporation, Japan). The Fluc and Rluc activities were measured using firefly luciferase and *Renilla* Luciferase Assay Reagents, E1500 and E2810 (Promega Corporation, USA), respectively according to manufacturer's instructions. The luciferase activity under the constitutive SV40 promoter was divided by their Rluc luciferase control activity to yield the normalized luciferase signals. Error bars indicate the standard deviation (n = 4).

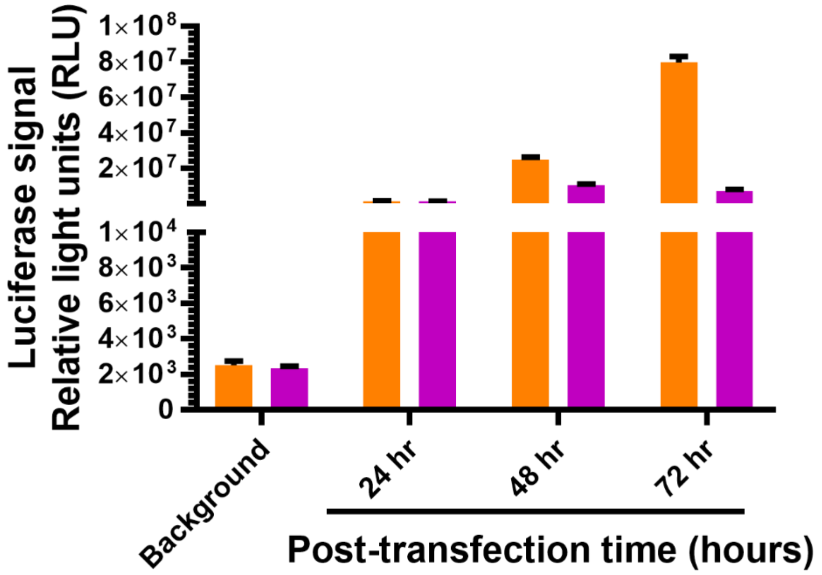


**Figure 4-figure supplement 1.** Effects of post-transfection period on FLUX signals. The pGL3 [*FLUX*/SV40] vector (orange) was co-transfected with the pRL-TK vector (purple, control vector) into HEK293T cells. Cells were collected at 24-, 48- and 72-hours post-transfection using Lux Lysis Reagent (LLR) and luciferase activities were independently measured. Non-transfected cells without any vector were monitored for their signals as background. Lux activities were monitored by adding a 100 µL of reagent cocktail consisting of 5 µM FMN, 100 µM HPA, 10 µM decanal and 100 µM NADH into a cell lysate which was freshly mixed with 50 mU of $C_1$ reductase. The luminescence signals were monitored for 10 sec with a 2-sec delay using an AB-2250 single tube luminometer (ATTO Corporation, Japan). The Rluc activity was measured using *Renilla* Luciferase Assay Reagent [E2810, Promega Corporation, USA) according to manufacturer's instructions. Error bars indicate the standard deviation (n = 4).

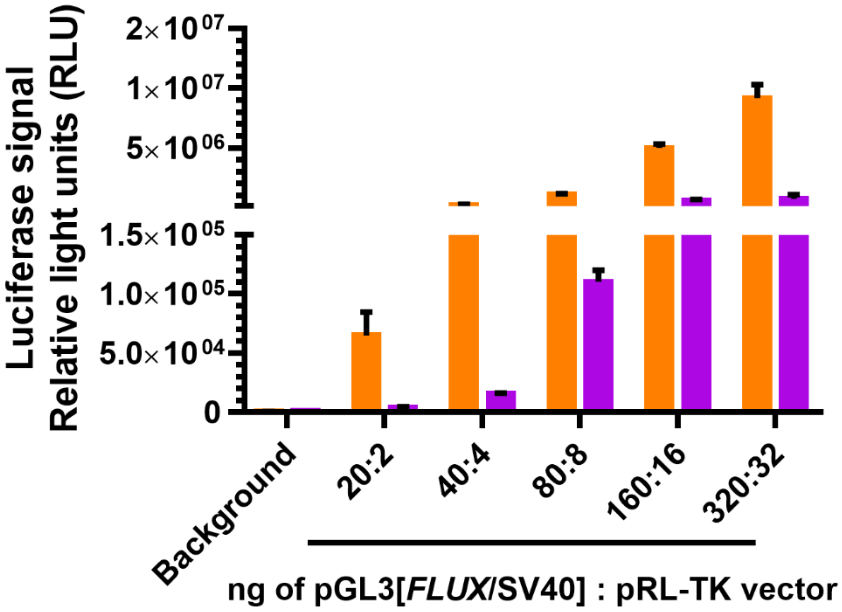


**Figure 4-figure supplement 2.** The FLUX signal (orange bar) from various amounts of pGL3 [*FLUX*/SV40] vector and their Rluc internal control signal (Purple bar). Two-fold seral dilutions of pGL3 [*FLUX*/SV40] vector from 320 to 20 ng were independent co-transfected with the pRL-TK vector as an internal control with a target:control vector ratio of 10:1 into HEK293T cells. The activity of Lux was measured by adding a 100 µL of a cocktail reagent consisting of 5 µM FMN, 100 µM HPA, 10 µM decanal and 100 µM NADH into cell lysate mixed with 50 mU of $C_1$ reductase. The Rluc activity was measured using *Renilla* Luciferase Assay Reagent [E2810, Promega Corporation, USA) according to the manufacturer's instructions. Luminescence signals were monitored for 10 sec with a 2-sec delay using an AB-2250 single tube luminometer (ATTO Corporation, Japan). Error bars indicate the standard deviation (n = 4).

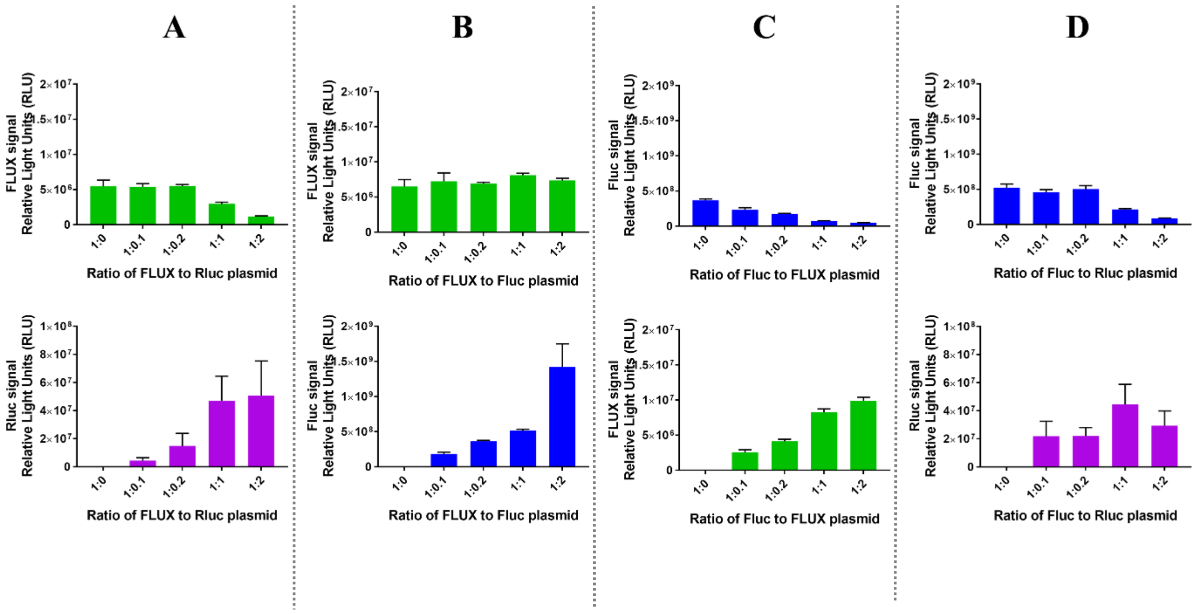


**Figure 5-figure supplement 1.** Individual signals of luciferase-based measurement at various ratios of targer:control vectors. Three types of luciferase vectors including pGL3[*luc*+/SV40] vector (blue), pGL3 [*FLUX*/SV40] vector (green) and pRL-TK vector (purple) were used as either target or control vectors for four combinations including **A.** FLUX/Rluc, **B.** FLUX/Fluc, **C.** Fluc/FLUX, and **D.** Fluc/RLuc. The target vector was co-transfected with the control vector at various ratios ranging from 1:0, 1:0.1, 1:0.2, 1:1, and 1:2 into HEK293T cells. Cells were collected at 48-hours post-transfection in either Passive Lysis Buffer or Lux Lysis Reagent and luciferase activities were independently measured. The activity of Lux was monitored by adding 100 µL of a cocktail reagent consisting of 5 µM FMN, 100 µM HPA, 10 µM decanal and 100 µM NADH into a cell lysate freshly mixed with 50 mU of $C_1$ reductase. The luminescence signal was monitored for 10 sec with a 2-sec delay using an AB-2250 single tube luminometer (ATTO Corporation, Japan). The Fluc activity was measured using firefly luciferase Assay Reagent (E1500, Promega Corporation) according to the manufacturer's instructions. Error bars indicate the standard deviation (n = 4).

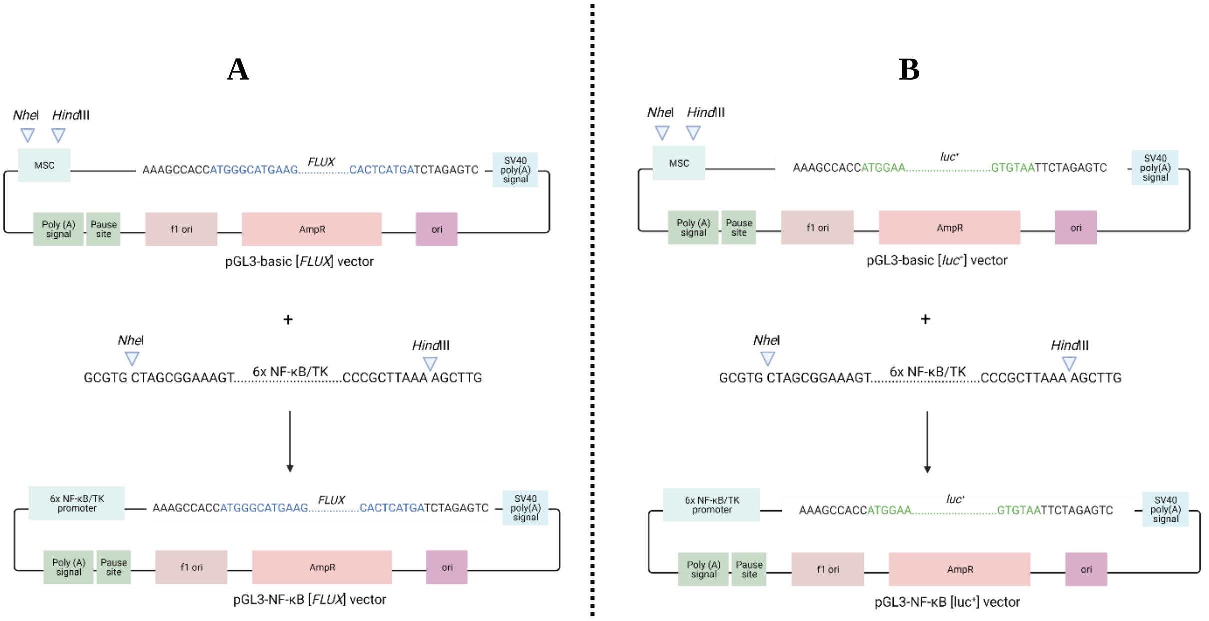


**Figure 6-figure supplement 1.** Maps of *FLUX* and *luc+* gene reporters under the control of six tandem repeats of the NF-κB transcriptional element with the TK promotor. **A.** Construction processes of the pGL3-NF-κB [*FLUX*/TK] vector and **B.** the pGL3- NF-κB [*luc+*/TK] vector. Construction processes of all vectors are similar. First, the pGL3 vector consisting of *FLUX* or *luc+* reporter gene was digested with *Nhe*I and *Hind*III restriction enzymes. Six tandem repeats of the NF-κB transcriptional element with TK promotor were amplified and digested by *Nhe*I and *Hind*III restriction enzymes. Then, the digested pGL3 vector was ligated with the digested FLUX gene to obtain the pGL3-NF-κB reporter gene/TK vector consisting of either *FLUX* or *luc+* as a reporter gene.
